# Gene loss during a transition to multicellularity

**DOI:** 10.1101/2021.02.16.431445

**Authors:** Berenice Jiménez-Marín, Jessica B. Rakijas, Antariksh Tyagi, Aakash Pandey, Erik R. Hanschen, Jaden Anderson, Matthew G. Heffel, Thomas G. Platt, Bradley J. S. C. Olson

## Abstract

Multicellular evolution is a major transition associated with momentous diversification of multiple lineages and increased developmental complexity. The volvocine algae comprise a valuable system for the study of this transition, as they span from unicellular to undifferentiated and differentiated multicellular morphologies despite their genomes being highly similar, suggesting multicellular evolution requires few genetic changes to undergo dramatic shifts in developmental complexity. Here, the evolutionary dynamics of six volvocine genomes were examined, where a gradual loss of genes was observed in parallel to the co-option of a few key genes. Protein complexes in the six species exhibited a high degree of novel interactions, suggesting that gene loss plays a role in evolutionary novelty. This finding was supported by gene network modeling, where gene loss outpaces gene gain in generating novel stable network states. These results suggest developmental complexity may be driven by gene loss, and not just by gene gain.

**Significance Statement:** Increased developmental complexity is thought to evolve mainly through genetic innovation and co-option. Comparison of the genomes of the closely related volvocines revealed that despite large changes in developmental complexity their genomes are exhibiting significant gene loss. Further, a burst of gene loss is shown to occur at the transition to undifferentiated multicellularity by gene inactivation and decay. This likely results in changes to protein-protein interactions within the cell, suggesting evolutionary novelty does not always require gene gain. Using empirical and modelling approaches we demonstrate gene loss can more easily produce protein network novelty than gene gain. We propose that gene loss can be a driver of biological innovation, allowing for reconfiguration of gene networks and differential use of existing functional repertoires.

## Introduction

The evolution of multicellular organisms is a prerequisite for the emergence of complex organismal body plans^1,2^. However, the genomic basis for multicellular evolution is poorly understood^3^. The volvocine algae are a valuable model system for studying multicellular evolution because they have undergone a recent transition to multicellularity (∼200 MYA)^4^, and the ∼100 member species span a wide range of developmental complexity within a size range from 10 µm to 3 mm^5^. This group includes unicellular *Chlamydomonas*, undifferentiated multicellular bowl-shaped *Gonium* and spheroidal *Pandorina* (8-16 cells), multicellular isogamous *Yamagishiella* (32 cells), multicellular anisogamous *Eudorina* (32 cells), and multicellular differentiated *Volvox* (>500 cells)^6,7^.

Based on morphology, differentiated multicellular volvocines were predicted to have evolved from their unicellular ancestors by stepwise acquisition of developmental processes, such as establishment of organismic polarity, genetic control of cell number, size expansion, and division of labor among cell lineages^6^ (Figure 1A, cartoons). However, phylogenetic approaches suggest that diversification among the volvocine algae was rapid^4^ and many of the genetic changes allowing for the evolution of developmental complexity took place early in the evolution of the volvocines. This suggests that genetic gains for each step might not fully explain the organismal complexity of the volvocine algae^8^. Indeed, the genomes of *Chlamydomonas reinhardtii, Gonium pectorale*, and *Volvox carteri*^*a*^ are highly similar and syntenic^8-10^. Moreover, a homolog of the cell cycle regulator and tumor suppressor *Retinoblastoma* (RB) is sufficient for undifferentiated multicellularity^8^, suggesting co-option of a developmental regulator can result in complex developmental patterns required for multicellularity^3^. Thus, the evolution and diversification of the volvocine algae is thought to have heavily relied on duplication, divergence and co-option of key functional elements.

**Figure 1.**
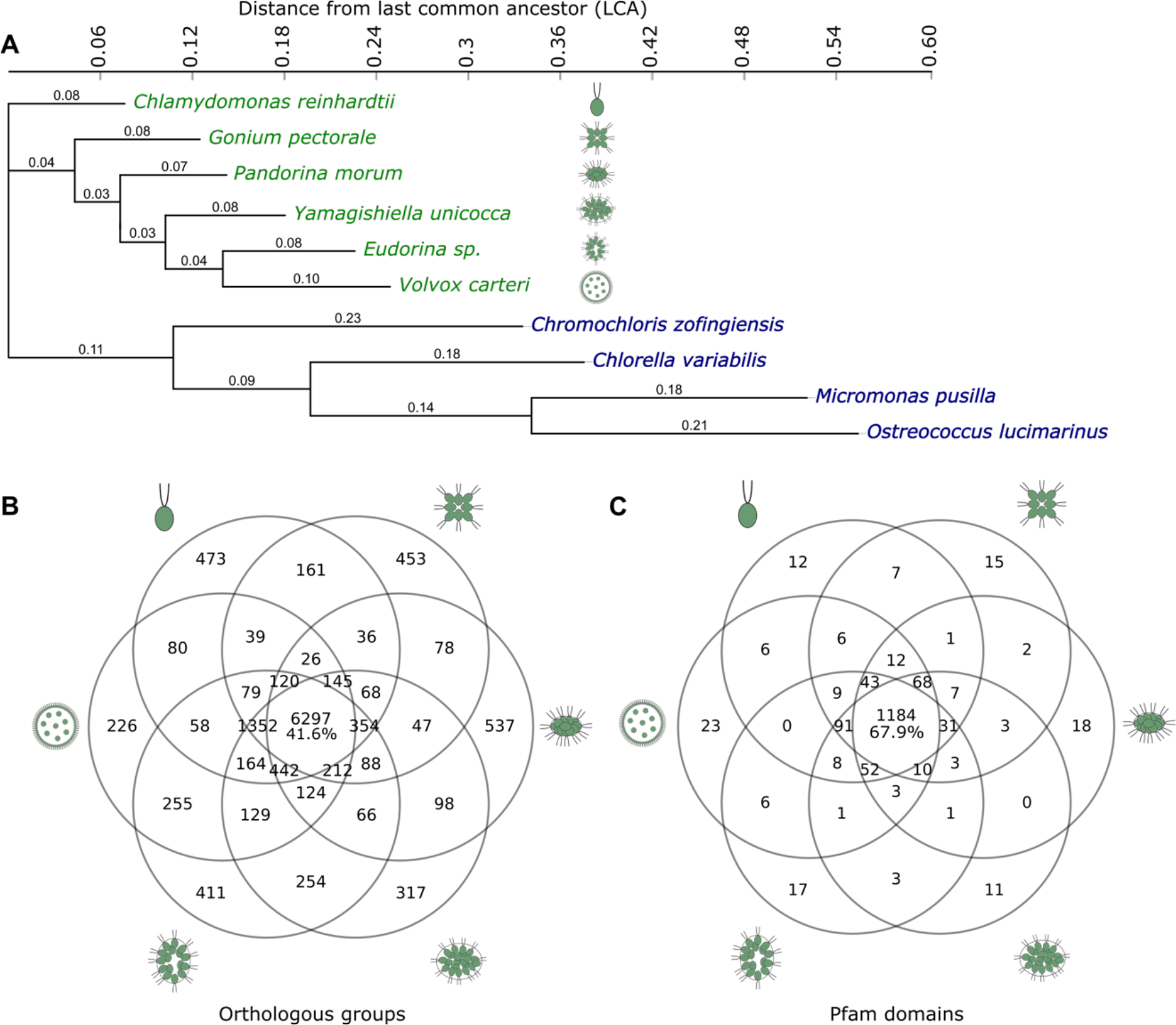
Volvocine algal genomes are highly similar. (**A**) Genome-wide phylogeny of the volvocine algae (green) compared to other chlorophycean outgroups (blue) inferred by PosiGene. Branch values indicate distance from the nearest node. Note *Chlamydomonas* is unicellular and closely related to colonial volvocines. (**B**) Orthologous groups shared between the six volvocine species (Supplementary File 1). (**C)**, Pfam domains shared between six volvocine species (Supplementary File 2).

Co-option, however, is unlikely to be the only evolutionary process behind volvocine multicellularity. Gene loss is found in many lineages undergoing evolutionary change^11-15^, and it is likely a major force in eukaryotic evolution^16^ with the potential to influence evolutionary trajectories^17^. Nevertheless, its action in concert with other evolutionary drivers is not well understood^18-21^. The study of gene loss is challenging because loss events can be the result of trait evolution (e.g. through relaxed selection), but they can also cause phenotypic adaptation^17^. Moreover, gene losses can impact an organism’s evolvability and speed of adaptation^22^, depending on their dispensability and position within the regulatory and protein-protein interaction network^16,22^.

To investigate the role of gene loss during the evolution of multicellularity, we report the assembly of *Pandorina morum* and annotation of genomes from *P. morum, Yamagishiella unicocca* and *Eudorina sp*.^a^ These newly analyzed genomes, in addition to those of *Chlamydomonas, Gonium*, and *Volvox*^8,9,23^, were subjected to comprehensive analysis of genic evolutionary trajectory. The high similarity of the genomes allows a detailed analysis evolutionary trajectory of genes as multicellularity complexity evolves. We demonstrate that significant loss of conserved genes has occurred in the volvocine lineage, and through a gene network model, we show that gene loss outperforms gene gain in generating stable novelty. Our results additionally suggest that only a few genes were co-opted for volvocine multicellularity. Furthermore, differential protein-protein interactions (PPI) are observed between species. These results suggest that gene loss at the transition to multicellularity likely impacts molecular network structure such that novel functions associated to changes in developmental complexity can evolve.

## Results

We compared the genomes of six closely related volvocine algae exhibiting the major changes in morphological complexity^6^ (Figure 1A cartoons, Supplementary Table S1) and validated their annotation using BUSCO^24^. BUSCO inferences suggest that all six annotated genomes are of high quality (Supplementary Table S2). Orthologous group and protein family domain (Pfams) catalogs for the analyzed species and their chlorophyte outgroups were compiled, and a genome-wide reconstruction of volvocine phylogeny was generated (Figure 1A) based on orthologous groups, which is in close agreement with previously determined phylogenies based on smaller gene sets^8,25,26^. Overall, 15,155 orthologous groups were identified; of these, 6,297 (41.6%) are shared among the six genomes, and 2,417 are species specific (15.9%, Figure 1B, Supplementary Table S3). Analysis of the evolutionary trajectory of Pfams^27^ in the six species shows 1,743 Pfam domains. Of these, 1,184 (67.9%) are shared in all six species, and 96 (5.5%) are species specific. Notably, 8,922 (58.9%) orthologous groups, and 1,479 (84.9%) Pfams are shared by at least 5 species (Figure 1C). These results support previous findings of the remarkable similarity found between the volvocine genomes^8,23^, despite their marked developmental and morphological differences.

Given the high degree of similarity between volvocine algae genomes (Figure 1B, C), we hypothesized that small differences between the *Chlamydomonas, Gonium, Pandorina, Yamagishiella, Eudorina* and *Volvox* genomes contribute to the evolution of multicellularity. Rates of evolution of shared genes and protein domains (7,942 orthologous groups and 1,338 Pfam domains) for the six species were analyzed by normalization to the distance from their Last Common Ancestor (LCA) to account for potential signatures of gene or domain gain and loss. This analysis revealed that 402 (5.1%) orthologous groups have undergone significant changes, with a bias towards gene loss (176 expanding orthologous groups, 226 contracting, Figure 2A outliers, shaded grey area). Surprisingly, while most Pfam domains were conserved, significantly expanding domains were less abundant than their contracting counterparts (51 and 127, respectively, Figure 2B outliers, shaded grey area). Pfam domain counts per gene did not show significant variation between the six species (Supplementary Figure S1), and contraction trends were not observed in highly conserved genes Actin, βTubulin, α-Tubulin, chloroplast 6erredoxin, mtATP-synthase α, mtATP-synthase β, and dynein heavy chain, (Supplementary Figure S2, Supplementary Table S4).

**Figure 2.**
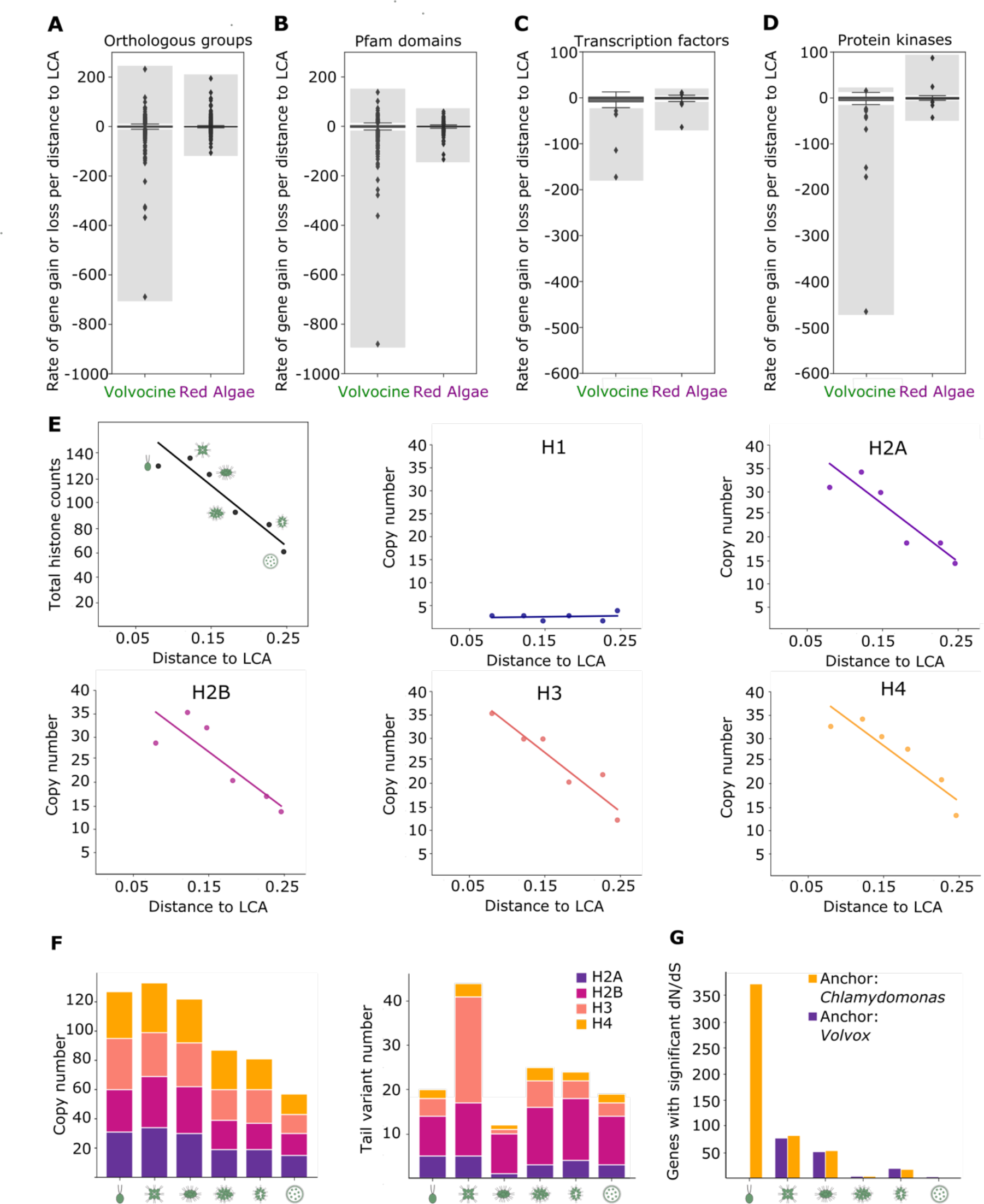
Gene loss outpaces gain in volvocine algae genomes. Distribution of the gene loss and gain rates per distance to LCA in the volvocine genomes compared to Rhodophyceae (red algae) of (**A**) orthologous groups, (**B)** Pfam domains, (**C**) transcription factors, and (**D**) protein kinases. Outlier groups, which represent significant loss (negative values) or gain (positive values) rates, are within the shaded grey area. (**E**) Histone copy number relative to distance to LCA shows a decreasing trend except for H1 in the volvocine genomes. Top left panel describes total histone counts per species. (**F**) Histone copy numbers (left) are reduced with minimal loss of tail variants (right). (**G**) Quantification of positively selected genes per species using *Chlamydomonas* (yellow) and *Volvox* (purple) as reference genomes.

Co-option of master regulators such as transcription factors (TFs) or protein kinases (PKs) is thought to be important for evolutionary innovation^8,28^. Analysis of all identified TF and PK families in the volvocine algae (47 and 69 respectively) revealed that 5 families of TFs (10.6%) underwent significant contraction, while none expanded significantly (Figure 2C, Supplementary Figure S3A). Similarly, 10 families of PKs (14.5%) underwent significant change, where 1 of the PKs expanded and 9 contracted (Figure 2D, Supplementary Figure S3B). These data suggest that volvocine algae have lost genes with conserved functions. Conversely, gene gain occurred at a much lower rate and frequency (Figure 2A-F, Supplementary Figure S3).

To gain further insight into gene loss as a mechanism of multicellular evolution, we analyzed histone genes for their evolutionary dynamics. Chromatin is an important regulator of gene expression, where post-translational modifications of histone N-termini are conserved and well understood^29^. Thus, histone genes were analyzed for their evolutionary dynamics. All families of histones, except linker H1, were reduced in number as developmental complexity increases in the volvocine algae (Figure 2E, Supplementary Table S5 – S6). Analysis of histone N-termini methylation site variation for H2A, H2B, H3, and H4 shows that H2A, H2B and H4 underwent copy number reduction (Figure 2F, Supplementary Table S6). However, Histone H3 exhibited greater N-terminal variant diversity in *Gonium* (Figure 2F, Supplementary Table S6). These data suggest that post-translational changes in histones may facilitate increases in developmental complexity through altering gene expression in the context of gene loss. Red algae (Rhodophyceae) also display various degrees of developmental complexity and exhibit signatures of gene loss^18^. Hence, their orthologous groups (Figure 2A), Pfam domains (Figure 2B), TFs (Figure 2C), PKs (Figure 2D) and histone (Supplementary Table S7) dynamics were examined in parallel to their volvocine counterparts. This analysis confirmed the presence of contraction signatures in red algae, agreeing with earlier reports^18^.

Few genes in the *Gonium* and *Volvox* genomes show a signature of undergoing positive selection, as measured by the ratio of non-synonymous to synonymous mutations (*d*N/*d*S)^8^. To assess the impact of positive selection on increases to developmental complexity, orthologous groups of genes with representatives in all six species were subjected to *d*N/*d*S analysis. Using *Chlamydomonas* as the reference for ratio calculation 77, 51, 4, 19 and 3 genes exhibited positive selection in *Gonium, Pandorina, Yamagishiella, Eudorina*, and *Volvox*, respectively (Figure 2G). Using *Volvox* as the reference, there were 372, 82, 53, 4 and 17 genes with a significant *d*N/*d*S value in *Chlamydomonas, Gonium, Pandorina, Yamagishiella*, and *Eudorina*, respectively (Figure 2G). Among multicellular volvocines, *Gonium* has a larger number of genes under positive selection, with the multicellular species examined having comparable albeit lower numbers of genes under positive selection. These results suggest that positive selection is not the only force involved in volvocine evolution; rather, it seems like processes of gene loss coupled to key co-option events might have set the stage for instances of positive selection, especially upon the transition to undifferentiated multicellularity.

The genomes of the volvocines are highly syntenic^8,23^, thus allowing the examination of the mechanism underlying gene loss for the 226 contracting orthologous groups at the sequence level (Figure 2A). Determination of the mechanisms behind gene loss at the sequence level is difficult to do in other phyla, where it is inferred indirectly through phylogenetic analysis^14,15^. The contracting groups contain 1,299 genes assembled into *Chlamydomonas* chromosomes, of which 1,184 (91.2%) in 199 (88.1%) orthologous groups had loci that were adequately assembled in all genomes for further analysis. This analysis demonstrated 43 of 1,184 genes in contracting orthologous groups are retained (preservation of orthologs across all species), while 1,141 loci exhibit gene loss (Figure 3A, B). The mechanisms of gene loss were then examined by tBLASTx to determine if remnants of the lost genes (gene decay) were present at the locus or if remnants of the gene were completely lost (gene deletion). Our results show gene decay is more abundant than deletion in all the colonial species; 72.7% of *Gonium*, 65.8% of *Pandorina*, 72.9% of *Yamagishiella*, 73.2% of *Eudorina* and 51.4% of *Volvox* gene losses show decay signatures (Figure 3A). Observed signatures of decay confirm that our observations of gene loss are not likely to be caused by technical issues with genome comparisons, but rather are, for the most part, evolutionary losses of gene function. Thus, the volvocines exhibit a burst of gene loss, primarily by decay during the transition to undifferentiated multicellualrity that continued as organismal complexity increased.

**Figure 3.**
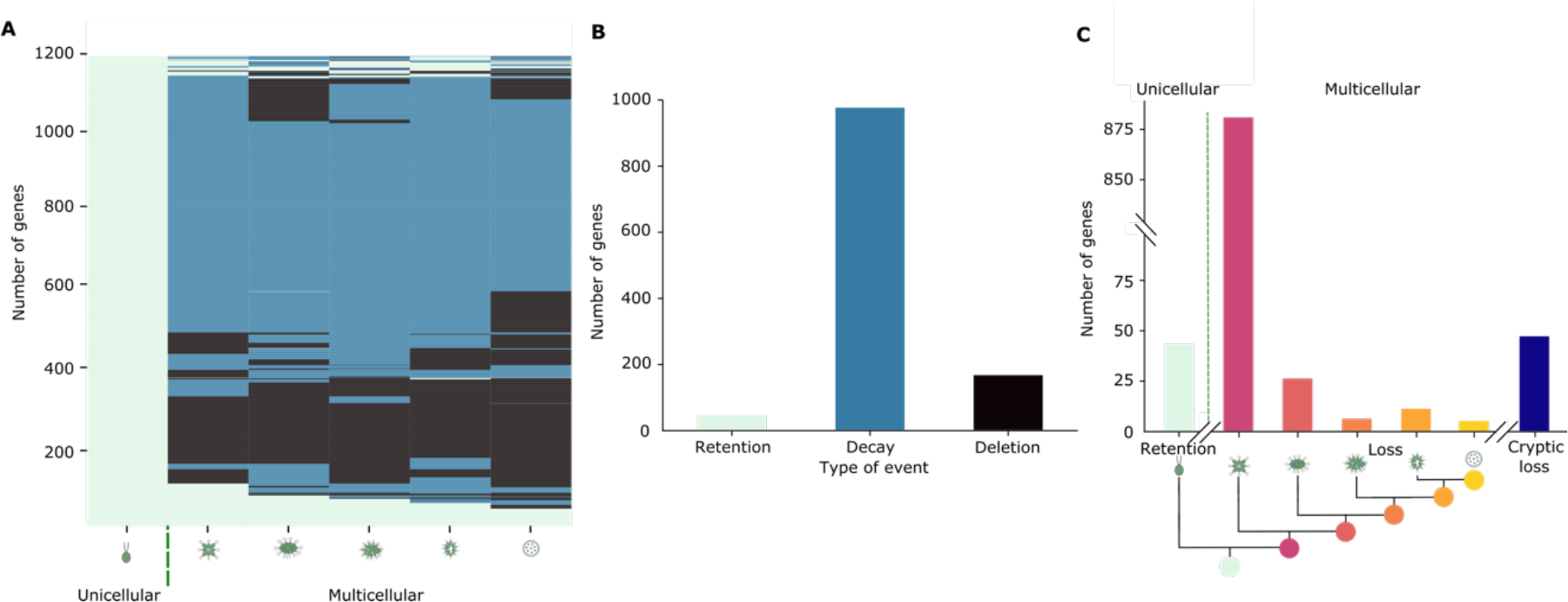
Gene loss in the volvocine algae occurs primarily by gradual decay. (**A**) Retention, decay, and deletion of 1,184 loci belonging to orthologous groups undergoing contraction in volvocine species. (**B**) Cross species quantification of retention (light green), decay (blue), and deletion (black) of loci in contracting orthologous groups. (**C**) Quantification of events of retention and loss by decay relative to evolutionary relationships between volvocine species. Losses that do not follow phylogenetic distributions are considered cryptic (dark blue).

Cross-species comparisons of retention, decay, and deletion events show that most candidate gene losses, 975 loci (82.3%), are due to decay (Figure 3B). Surprisingly, our analysis also suggests 1,017 (85.9%) loci are gene loss candidates in all five multicellular species. Of these, 880 (74.3%) occurred by gradual decay (Figure 3C, Supplementary Table S8). The presence of decay signatures eliminates the possibility that these loci represent gene gains of *Chlamydomonas*. In the case of deletions, only 166 loci (14.0%) were completely absent in all multicellular species, which could either represent complete gene loss in the multicellular lineage or *Chlamydomonas-*specific gene gains. GO-term analysis of *Chlamydomonas* orthologs of retained (24 of 43 genes), decayed (234 of 975 genes) and deleted genes (42 of 166 genes) was also performed. Surprisingly, genes associated to conserved biological processes (e.g. metabolic processes, ion transport) and molecular functions (e.g. protein kinase activity, nucleic acid binding) have undergone loss by decay, deletion, or both (Supplementary Figure S4).The predominance and distribution of decay signatures suggest that a major gene loss event(s) occurred after the LCA of colonial volvocine algae diverged from the lineage of rather than that the *Chlamydomonas* lineage experienced extensive gene gains after diverging from multicellular volvocines.

Previously, genes important for *Volvox* development were hypothesized to involved in the evolution of multicellularity^3,8,23^. used phylogenetic analysis to gain deeper understanding of co-option in the volvocine algae. Cyclin D1 genes (Supplementary Figure S5F), certain extracellular matrix (ECM) related genes (Supplementary Table S9, Supplementary Figure S6, S7), and the *Volvox* cell differentiation gene *regA*^*30*^ follow a duplication and divergence pattern that coincides with increases in complexity. However, other developmentally relevant genes such as other cyclins (Supplementary Figure S5 A-E), *invA, B, C*, (Supplementary Figure S8) and *glsA*, (Supplementary Figure S9) do not. Genes that encode intrinsically disordered proteins (IDPs) were also examined, since these proteins might be co-opted for novel developmental functions in the volvocine algae^31,32^. However, no significant change in the IDP content, frequency or distribution was observed between the volvocine algae genomes (Supplementary Figure S10).

To examine the role of genetic innovation in volvocine evolutionary history, their genes were assigned to nine phylostrata (Supplementary Figure S11). This analysis revealed that 51.1 to 69.8% of the genes per species fell under phylostratum 1 (PS-1, cellular organisms) and PS-2 (Eukaryotes) (Supplementary Figure S11). Conversely, 0 to 4.3% of the genes per species were found in PS-7 (family). These results suggest that co-option and expansion of key genes is a relevant, but does not coincide with stepwise increases in developmental complexity as previously thought^6^.

To understand how developmental complexity could evolve in the context of significant gene loss in the volvocine algae, the dynamics of their protein-protein interactions (PPIs) were examined. We reasoned that changes in PPIs could lead to impactful changes in volvocine developmental complexity, even though few genes are undergoing positive selection or co-option (Figure 2G). Examination of silver-stained 2D-PAGE gels of total protein extracted from *Chlamydomonas, Gonium* and *Eudorina* shows extensive differences in the proteomic makeup of each species (Figure 4A, Supplementary Figure S12).

**Figure 4.**
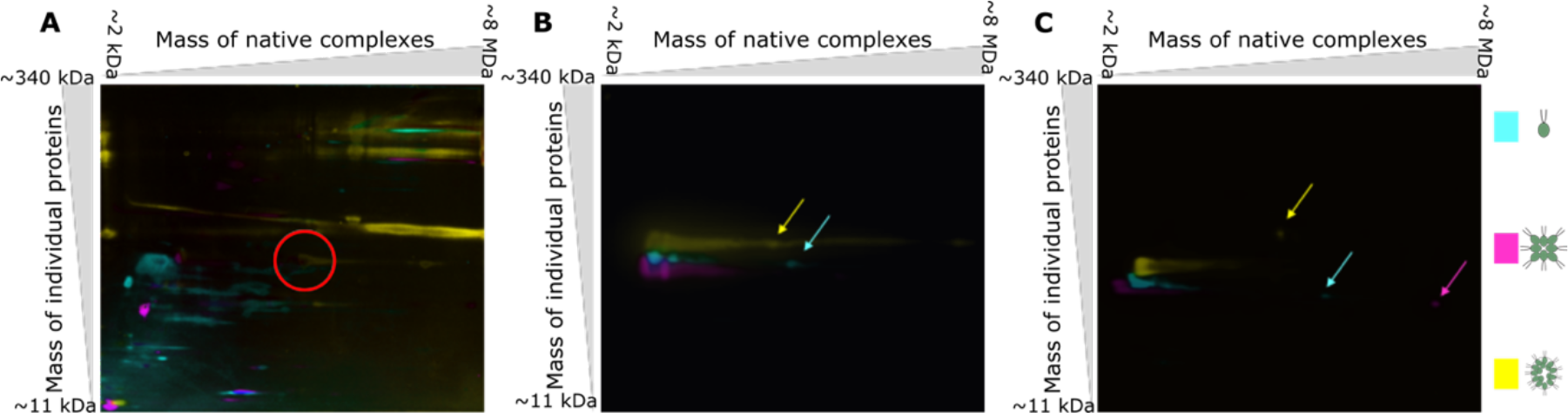
Protein-protein interactions differ among volvocine species. Blue Native (BN) PAGE followed by SDS-PAGE of lysates from *Chlamydomonas* (pseudocolored cyan), *Gonium* (pseudocolored magenta), and *Eudorina* (pseudocolored yellow). (**A**) Overlay of silver stains. RuBisCO signal is circled in red. (**B**) Overlay of western blot signals for α-tubulin. Tubulin signals for species-specific complexes are indicated by color coded arrows. (**C**) Overlay of western blot signals for β-actin. Actin signals from unique complexes are indicated by color coded arrows.

Changes in morphological and developmental complexity in the volvocine algae are thought to be related to the cytoskeleton^33^. This is because cytoskeletal elements common to *Chlamydomonas* and *Volvox* are involved in cell division in both species, and in the latter also in inversion^33^. Hence, PPIs of the actin and tubulin cytoskeleton were examined. Tubulin and actin proteins are highly conserved in the volvocine algae (Supplementary Figure S13), and as expected, complexes of similar size were found in *Chlamydomonas, Gonium*, and *Eudorina* 2D-PAGE immunoblots (Figure 4B, C, Supplementary Figure S12). Minor differences in migration patterns for both proteins in the denaturing dimension are likely caused by post-translational modifications. Notably, novel α-tubulin complexes were shifted along the native axis, suggesting the presence of larger, species-specific complexes containing tubulin (Figure 4B, arrows). Similarly, 2D-PAGE immunoblots for β-actin showed a common streak in all three species, but also displayed species-specific actin-containing complexes that are larger than the shared complex (Figure 4C, arrows). Cultures used to extract protein were asynchronous; hence, the differential migration patterns between species are not likely to result from comparing species at differential life stages. Taken together, these results indicate that while the genomes of the volvocine algae are highly similar and experiencing gene loss, PPIs between conserved gene products are forming species-specific complexes, which could account for the observed differences in biological complexity between the volvocines.

To evaluate the potential of gene loss as a mechanism of evolutionary transitions, gene loss and gain were modeled using an adaptation of the Wagner gene regulatory network model^34^. Our model mirrors the effects of gene gain in Wagner’s original model, which demonstrated that networks more often reach a new equilibrium state following an intermediate number of gene duplications than when there are many or few duplications^34^. Our model predicts similar outcomes following gene loss with the proportion of novel stable gene network configurations being higher following gene deletions as compared to an equal number of duplications (Figure 5). Upon iterating the model over a broad range of network connectivity, similar results were obtained (Supplementary Figure S14). Through network-wide loss of robustness, novel stable network states might result in regulatory rewiring that allows for major changes to developmental programs. In sum, our evidence indicates that the signature of gene loss in the volvocine genomes may represent an example of a broadly used mechanism for the evolution of developmental complexity.

**Figure 5.**
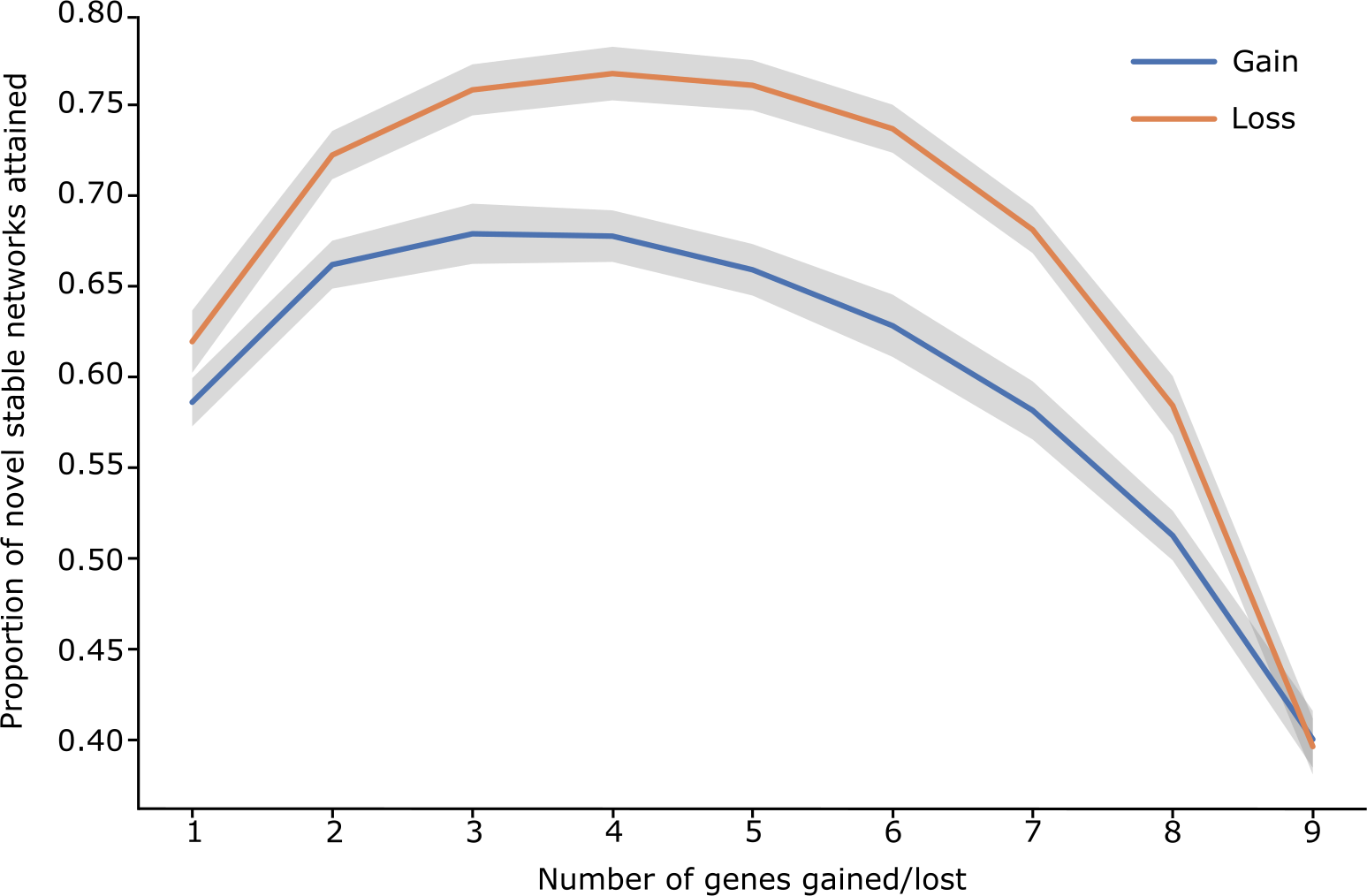
Gene loss in a fully connected network (*c*=1) of genes (*N*=10) yields a higher proportion of novel stable network states than gene gain. Wagner model simulates loss (orange) and duplication (blue) of *k* genes. Grey shading represents variance.

## Discussion

Our work challenges the common assumption that biological and genomic complexity are positively correlated^35,36^. If this were the case, the developmental gains in *Volvox* and species that exhibit seemingly intermediate steps of developmental complexity (typified by *Gonium, Pandorina, Yamagishiella* and *Eudorina*) might be the consequence of sequential acquisition of novel genes and/or the result of co-option events correlating to each developmental gain^3,30,37,38^. However, the six analyzed genomes did not show evidence of extensive gene duplication, subspecialization or neofunctionalization. Rather, we discovered that a very small set of expanding and co-opted genes are found in the five multicellular species (Supplementary Table S9, Supplementary Figure S5 – S9), some of which have an evolutionary signature of sequential co-option. With few exceptions (e.g., *reg*, extracellular matrix (ECM) genes, cyclin D), it was not possible to assign novel genes to specific developmental gains in the volvocine algae, despite the complexity of these novel traits.

Notwithstanding the obvious similarities between volvocine genomes (Figure 1B, C), genetic losses in conserved genomic regions have occurred in volvocine history (Figure 3). Orthologous groups and Pfam domains both show more numerous significant gene losses than they do gains, and the rate of loss for some of these groups can be quite dramatic (Figure 2A, B), even in the absence of significant variation in the number of Pfam domain counts per gene between the six species (Supplementary Figure S2). These results agree with previous work describing pervasive Pfam loss events across eukaryotes ^39^. Regulatory and histone genes, which are important for development ^8,28,40^, are not exceptions to this trend (Figure 2C, D, E). The non-uniform distribution of gene loss events across the five multicellular species (Figure 3A, C), as well as the signatures of decay found for many of these genes (Figure 3B), contend that the observed patterns are not a consequence of annotation disparities or analytical biases. Events of gene loss throughout evolution are not unique to the volvocines, suggesting that this is a widespread phenomenon^18-20^. However, this is the first time to our knowledge that the mode of gene loss has been traced within a lineage. The mechanisms underlying the observed rates of decay for genes lost remain enigmatic, but their presence allows for the inference that most gene loss events are *bona fide* and, not *Chlamydomonas* specific gene gains. Evidence for gene loss through gradual decay in the context minimal duplication and divergence differs from the model of differential loss of paralogs that has been proposed for metazoans ^14^. It also appears to differ from other lineages^39^ in the sense that there is no clear functional bias related to gene loss (Supplementary Figure S4); instead, it would seem that a wide array of functions are differentially preserved. This supports the possibility that there is an adaptive role for gene loss in the context of phenotypic innovation and developmental rewiring.

Previous reports posited that gene duplication is a costly process due to the expense of synthesizing the gene products and the impact of increased function/gene dosage alteration on the cell^41^. Moreover, novel stable regulatory networks can be readily generated via gene loss (Figure 5, Supplementary Figure S14). Hence, both theoretical and biological systems suggest that gene gains are neither the only nor the fastest route to biological innovation. Our data supports the idea that the transition to multicellularity is driven by re-purposing existing genes for new functions while eliminating others as opposed to by generating of extensive genetic novelty.

The patterns of gene loss and PPI diversification in the volvocine algae illustrate vigorous genomic and proteomic shifts that likely underlie changes to evolutionary and developmental constraints between members of a single lineage. A significant proportion of the gene losses (∼55%) identified in the volvocine algae were shared by the four colonial species, whereas other gene loss patterns were less numerous (Figure 3C). The ancestor of undifferentiated *Gonium* (and the other multicellular species studied) would then appear to be the hotspot for initiation of gene loss and co-option. Analysis of genes with a signature for positive selection shows *Gonium* has the largest number of genes under positive selection among the species we studied (Figure 2G), and it appears to have undergone diversification of histone H3 N-terminal tails (Figure 2F). Perhaps the developmental complexity portrayed by *Volvox* was made possible by the same mechanisms and tools visible in *Gonium*, such as co-option of RB^3^. Although the volvocine transition to undifferentiated multicellularity is a single event, differentiated multicellularity has arisen multiple times^11,42^. The volvocine spherical colony morphology has also evolved at least twice^43^; *Astrephomene* is notably similar to *Eudorina* phenotypically, it is closely related to *Gonium* (Goniaceae) and its embryogenesis and development is markedly different from that of other spheroidal volvocines^43^. These examples support the possibility that complex multicellularity is a relatively straightforward consequence of simple multicellularity. Should this be the case, a future challenge would be to test this hypothesis in other multicellularity models as means to assess if the true evolutionary transition lies in establishing the conditions that stabilize undifferentiated multicellularity through loss of gene functions that might be necessary to maintain the unicellular state. In sum, our results indicate that biological innovation coincides with gene loss events, which could allow for reconfiguration of gene networks, differential use of existing functional repertoires, and reduction of the impact of genetic gains.

## Materials and Methods

### Strains and genomes

*Chlamydomonas reinhardtii* CC-503 MT^+^ genome version 5.5 ^9^(GCA_000002595.3), *Gonium pectorale* K3-F3-4 MT^-^ NIES-2863 ^8^(GCA_001584585.1), and *Volvox carteri f. nagariensis*, Eve S1^23^ genome versions 1 (GCA_000143455.1) and 2.1 (available on Phytozome)^23,44^ were used for all analyses. Genome assemblies for *Yamagishiella* and *Eudorina* were kindly provided by Dr. Hisayoshi Nozaki before publication^45^. All analyses involving *Yamagishiella* were performed using *Y. unicocca* strains 2012-1026-YU-F2-6 MT^+^ NIES-3982, and 2012-1026-YU-F2-1 MT^-^ NIES-3983. For *Eudorina, Eudorina* sp. strains 2010-623-F1-E4 female NIES-3984 and 2010-623-F1-E2 male NIES-3985 were utilized. Analyzed red algae genomes have been published previously: *Porphyra umbilicalis*^46^ (GCA_002049455.2), *Pyropia yezoensis* ^47^ (GCA_009829735.1), *Chondrus crispus*^48^ (GCA_000350225.2), *Porphyridium purpureum*^49^ (GCA_000397085.1) and *Cyanidioschyzon merolae*^50^ (GCA_000091205.1). Chlorophyte outgroup species genomes have been published previously: *Chromochloris zogingiensis* (available on Phytozome)^44,51^, *Chlorella variabilis* ^52^(ADIC00000000), *Micromonas pusilla* ^53^(GCA_000151265.1), and *Ostreococcus lucimarinus*^54^ (GCA_000092065.1).

### Algae growth conditions and library preparation

Cultures of *Pandorina morum* 5, a gift from Dr. Hisayoshi Nozaki, were initially decontaminated by collecting colonies grown in Standard Volvox Media (SVM) by centrifugation at 1000 *g* for 5 min at 25°C and removal of the growth media. These colony pellets were overlaid with 50% Percoll® (Millipore-Sigma) in SVM plus Acetate (SVMA) followed by illumination with 100 uE of light at the top of the tube. The top 1 mL of colonies that were able to phototax through this were then plated for single colonies on SVMA agar plates supplemented with 50 ug/mL carbenicillin, 50 ug/mL cefotaxime, 10 ug/mL trimethoprim and 20 ug/mL chloramphenicol. Cultures from single isolates were established in SVM and grown with 0.5% CO_2_ bubbling at 30°C, 350 µmol m^-1^sec^-1^ light intensity on a 14:10 day:night cycle. Cultures were harvested by centrifugation at 600*g* for 10 min with 0.005% Tween-20. DNA was isolated using a magnetic bead protocol to obtain high molecular weight DNA^55^. Illumina TruSeq PCR-free libraries were prepared from DNA that was either unfractionated or fractionated on a Pippin Prep (Sage Science) in the size ranges of 1kb, 2kb, 5kb and 10kb. Following library preparation, they were sequenced on an Illumina HiSeq2500 in Rapid Run mode to an estimated coverage of 25X. High molecular weight DNA was also subjected to Pippin Prep fractionation for 15 kb molecules, followed by PacBio SMRTbell library preparation. Libraries were sequenced on a PacBio Sequel system to an estimated 10X coverage. All sequencing was performed by Genewiz (South Plainfield, New Jersey, USA).

*Yamagishiella unicocca* 2012-1026-YU-F2-1 (NIES-3983, MT^-^) and *Eudorina* sp. strain 2010-623-F1-E4 (NIES-3984, female) were axenically cultured for RNA-seq experiments. Algae were grown SVM with 0.5% CO_2_ bubbling at 30°C, 350 µmol m^-1^sec^-1^ light intensity on a 14:10 day:night cycle. Triplicate cultures were harvested by centrifugation at 600*g* for 10 min with 0.005% Tween-20. RNA was extracted using a Quick-RNA Plant Miniprep Kit (Zymo Research). RNA-Seq libraries were prepared using the TruSeq Stranded mRNA kit (Illumina, Inc. San Diego, California, 20020595) according to manufacturer’s conditions.

Axenic cultures for 2D PAGE were grown asynchronously as follows: *Chlamydomonas reinhardtii* strain 21*gr* (*Chlamydomonas* collection CC-4889) was grown in TAP media to a density of ∼10^6^ cells/mL in ambient conditions with 0.5% CO_2_ bubbling. *Gonium pectorale* (strain Kaneko4, MT^+^ NIES-1711) was grown in SVM + acetate media to a density of ∼10^6^ colonies/mL in ambient conditions with 0.5% CO_2_ bubbling. *Eudorina* sp. strain 2010-623-F1-E4 (NIES-3984, female) was grown in SVM to a density of ∼10^6^ colonies/mL at 30°C with 350 μmol m^-1^ sec^-1^ light intensity on a 14:10 day: night cycle with 0.5% CO_2_ bubbling.

### Genome assembly

Illumina reads were assembled *de novo* with ABySS 2.1.5^56^ using a range of k-mers from 21-89, in increments of 4. Unitigs from each of the assemblies were then filtered against a database of known bacterial genomes to remove presumed bacterial endophyte sequences. Remaining unitigs were assembled with ABySS under the same conditions as above. Resulting scaffolds were merged to PacBio reads and assembled using Canu 1.8^57^ (Bioproject PRJNA787305).

### Evidence-based gene prediction

The gene models for *Yamagishiella* and *Eudorina* genomes were generated using RNA-seq data and the AUGUSTUS 3.2.3 software package^58^, in a manner similar to gene predictions for *Volvox* and *Gonium*^8,23^. The gene models for *Yamagishiella* and *Eudorina* genomes were generated using RNA-seq data (Bioproject PRJNA787306 and PRJNA787302) and the AUGUSTUS 3.2.3 software package^58^, in a manner similar to gene predictions for *Volvox* and *Gonium*^8,23,59^. AUGUSTUS predicted genes were merged with the genome mapped reads and de-duplicated to remove any splice variants.

### Identification of orthologous groups

Orthologous groups of volvocine algae and red algae genomes were determined using OrthoMCL 2.0.9^60^. The optimum inflation value was determined empirically by testing a variety of values ranging from 1.2 to 4.0 in increments of 0.1 (Supplementary Figure S15). The inflation value of 1.9 was used for all analyses (see Supplementary File 1).

### Identification of Pfam domains

The diversity and abundance of known Pfam domains in the volvocine algae and red algae genomes were identified using using Pfam Scan^61^ and database version 31. Pfam Scan and associated data collection Perl scripts available at: ftp://ftp.ebi.ac.uk/pub/databases/Pfam/Tools/ ^27^(see Supplementary File 2).

### Analysis of other genomic features

Transcription factors and protein kinases in the annotated volvocine algae genomes were identified using the iTAK version 1.7 program and database^62^. The *Chlamydomonas* gene definitions on GenBank were used to identify histone H1 (Accession #XM_001696120) and quantified (Figure 2E second panel, Supplementary Table S5). Histones H2A, H2B, H3, H4 (Accession # L41841) genes in the volvocine algae were identified using the reciprocal best BLAST hit method with a e-value cutoff of 1e-5. H2A, H2B, H3 and H4 genes were further filtered using e-values of 1e-10 and quantified (Figure 2E panels 3-6, Figure 2F first panel, Supplementary Supplementary Table S6). The first 30 amino acids from each histone were used for analysis of duplication and presence of predicted post-translational modification sites (lysine methylation) using Methylsight^63^. N-tails per histone per species that had different predicted sites with treshold >=0.5 were considered tail variants and quantified (Figure 2F second panel, Supplementary Table S6). Histone genes were also identified in the five red algae genomes using gene definitions from the *Chondrus* genome annotation (Supplementary Table S7). Eight internal control genes, Actin, mitochondrial ATP-sythase α subunit (mATP-A), mitochondrial ATP-sythase β subunit mATP-B, α-tubulin, β-tubulin, β-2 tubulin, flagellar dynein heavy chain, chloroplast ferridoxin were identified using OrthoMCL (Supplementary Table S4).

### Identification of gene count trends

Rate of gene number change was calculated by applying a linear regression model on the relationship between gene counts per orthologous group per species and distance of said species from their Last Common Ancestor (LCA, Figure 1A). Only groups with non-identical gene copy numbers and comprising genes from at least 2 species were used for this analysis. Rate of change values per orthologous group were subjected to quartile analysis to identify outlier groups (Figure 2A). This same analysis was performed on Pfam domains (Figure 2B), transcription factors (Figure 2C), and protein kinases (Figure 2D) for volvocine species and for red algae species. Regression and quartile analyses were performed using custom Python scripts.

### Identification of positively selected genes

Positively selected genes were identified using the PosiGene pipeline^64^. Two independent PosiGene analyses were performed using *Chlamydomonas* and *Volvox* genome annotations as anchors. Chlorophytes *Chlorella variabilis, Micromonas pusilla, Ostreococcus lucimarinus* and *Chromochloris zofingiensis* were used as outgroups. Similarly, the red algae genes under positive selection were identified using *Porphyra umbilicalis* as anchor species as it has previously been reported to be closest to the LCA of red algae^65,66^. A gene was considered to be positively selected if the *d*N/*d*S value (HA foreground omega) was greater than 1 and the Bonferroni-corrected p-value was less than 0.05. As a part of the workflow, PosiGene generates a phylogenetic tree from the isoform assignments and calculates the distance to LCA (Figure 1A, Supplementary Figure S16).

### Characterization of gene loss patterns

To evaluate annotation completeness, BUSCO v4.0.2^24^ was run in protein mode using the Chlorophyta lineage dataset for each volvocine species (Supplemnentary Table S2).

Orthologous groups showing significant contraction (Figure 2A) were selected for gene loss analysis based on whether they included *Chlamydomonas* genes. *Chlamydomonas* loci that were not assigned to chromosomes were not analyzed further. Orthologous groups that included *Chlamydomonas* loci with the aforementioned characteristics (target loci) served as anchors for obtention of genes up and downstream, called the ‘syntenic neighborhood’. Orthologs for loci in the syntenic neighborhood, as well as target loci for *Gonium, Pandorina, Yamagishiella, Eudorina* and *Volvox* were retrieved from their respective orthologous groups. Instances of no ortholog found for target loci were subjected to tBLASTx^67^ 2.2.18+ searches using tailored databases based on syntenic neighborhoods for each species. Non-informative (loss of syntenic neighborhoods or no BLAST results) genes were omitted from further analyses. tBLASTx hits were filtered for quality (e-value < 1e-5). Hits passing quality threshold were considered to be evidence of a gene remnant suggestive of loss via decay. Genes with no hits passing quality threshold and conserving synteny are candidate deleted genes. Decayed and deleted genes were assigned to bins corresponding to their pattern of absence (e.g. a gene lost by whichever mechanism only in *Gonium*, but present in the other clades; see Figure 3C and Supplementary Table S8), or by type of event, where decayed gene clusters consist of orthologs with at least one signature of decay in multicellular species, and deleted gene clusters are those where all losses happen via deletion (Figure 3B). Percentages of decay and deletion cases were calculated, and a Chi square goodness-of-fit test was applied to determine statistical significance of observed patterns using a uniform distribution as expected probability distribution. All analyses except for tBLASTx search were performed using custom Python scripts. Treemaps for GO analysis of retained, decayed and deleted genes were generated using REVIGO^68^ with default parameters and against *Chlamydomonas reinhardtii* database.

### Identification and analysis of known co-opted genes

Cell cycle genes in *Pandorina, Yamagishiella* and *Eudorina* genomes were annotated as previously described^8,23^ using tBLASTn. Transcriptome data and manual curation^8,23^ was used to prepare gene models for *Yamagishiella* and *Eudorina*. Known pherophorins are composed of two pherophorin domains (Pfam DUF3707) that are connected by a variable length hydroxyproline-rich repeat region^69^. All genes containing two or more pherophorin domains were identified as pherophorin genes. Matrix metalloproteases (MMPs) contain a single metalloprotease domain (Peptidase M11, PF05548). Genes containing this Pfam domain were selected for the existence of the [HQ]EXXHXXGXXH motif in the gene model^70^. The *invA, invB, invC* and *glsA* genes were identified in *Pandorina, Yamagishiella* and *Eudorina* by the presence of kinesin (PF00225), TPT (PF03151), glycosyl-transferase for dystroclygan (PF13896), and DnaJ (PF00226) domains respectively^37,38^.

### Prediction of lineage specific genes

Phylostratigraphy^71^ was used to predict the lineage of genes in the volvocine algae genomes as described previously^8^. The phylogenetic classes of volvocine algae were defined in the input data as per NCBI taxonomy. All protein sequences were searched against the NCBI non-redundant database from version 2.6+^67^, with an e-value cutoff of 1e-3. Phylostratigraphic classification of each gene ranged from PS-1 to PS-9. Gene level phylotratigraphic classification was converted to Pfam domain level classification.Phylostratigraphy^71^ was used to predict the lineage of genes in the volvocine algae genomes as described previously^8^. The phylogenetic classes of volvocine algae were defined in the input data as per NCBI taxonomy. All protein sequences were searched against the NCBI non-redundant database from version 2.6+^67^, with an e-value cutoff of 1e-3. Phylostratigraphic classification of each gene ranged from PS-1 to PS-9. Gene level phylotratigraphic classification was converted to Pfam domain level classification.

### Identification of protein disorder and protein binding site disorder

Protein disorder was calculated using DISOPRED3^72,73^, that yields percentage disorder in each protein sequence as well as that of protein binding sites. The frequency distribution of percentage protein disorder was calculated and plotted.

### 2D gel electrophoresis and silver stain

Approximately 10^6^ cells/colonies of *Chlamydomonas, Gonium*, and *Eudorina* were pelleted and snap frozen in liquid nitrogen. Cells were lysed by six rounds of liquid nitrogen freeze/thaw lysis in in low salt TBS (50 mM Tris, 50 mM NaCl, pH 7.5) and protease/phosphatase inhibitors (Plant Protease Inhibitor Cocktail (Sigma, Burlington, Massachusetts), 1 mM Benzamidine, 1 mM PMSF, 1 mM Na_3_VO_4_, 5 mM NaF, 0.5 mM β-glycerophosphate, 0.1 mM EDTA)^74^. Insoluble material was pelleted by centrifugation at 20,000 **g** for 10 min at 4°C. Supernatant was removed and protein content was quantified by Bradford assay (BioRad, Hercules, California)^75^. For blue native PAGE, 20 µg of protein per well was loaded into a 4-12% native PAGE gel (Fisher, Waltham, Massachusetts) with native running buffer (Fisher) and light blue buffer at the cathode (Fisher). Gel was run until the Coomassie dye front had exited the gel. Lanes were excised using a razor and incubated in 1x Tris-MOPS running buffer (Genscript, Piscataway, New Jersey) with 100 mM DTT and 5 mM sodium bisulfite at 55°C for 10 min. Gel slice was loaded into a NuPAGE 4-12% Bis-Tris gel (Fisher) and run with Tris-MOPS running buffer with 5 mM sodium bisulfite in the cathode buffer until the blue dye front had exited the gel. Silver staining was carried out using a Silver Staining Kit (Pierce) according to manufacturer’s protocols.

### Western blot and antibodies

Western transfer was performed in Tris-Glycine transfer buffer to a nitrocellulose membrane (0.45 µm pore, Pall Scientific, Port Washington, New York)^74^. Membranes were blocked in 1% milk (α-tubulin) or 5% BSA (β-Actin) for 1 hr. Primary antibodies (mouse anti-α tubulin, Sigma T6074 1:5000; mouse anti-β actin HRP, Santa Cruz sc-47778 HRP, 1:5000, Dallas, Texas) were incubated at 4°C overnight with gentle shaking. Membranes were washed with TBS 0.1% Tween-20 (TBST) and tubulin blots were probed with anti-mouse secondary (Pierce 31430, Rockford, Illinois) for 1 hr. Actin blots did not require a secondary antibody, as mouse anti-β actin is already HRP conjugated. Membranes were developed using Premium Western Blotting Reagent (LI-COR, Lincoln, Nebraska) and scanned for chemiluminescence on a C-Digit Scanner (LI-COR).

### Phylogenetic analysis

In order to determine phylogenetic relationships, the genes from all six volvocine genomes and *Coccomyxa subellipsoidea* (Chlorophyta, GCF_000258705.1) were aligned using MUSCLE v3.8.425 ^76^ and a phylogenetic tree was produced using RaxML v8^77^ with protein gamma model and automatic model selection. A rapid bootstrapping analysis with 1000 bootstrap replicates was performed and a majority ruled consensus tree was produced.

### Regulatory network model

A Wagner regulatory network model^34^ was designed as follows: a gene network of *N* genes whose on or off expression state at a time *t* is given as:

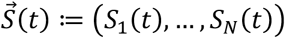

Where *S*_*i*_(*t*) is the expression state of the *i*^*th*^ gene at time *t, S*_*i*_*=*0 reflects an off state, and *S*_*i*_*=*1 reflects an on state. The state *S*_*i*_(*t*) is determined by the network of interactions affecting gene *i*. The expression state of gene *i* can be changed due to regulatory interactions. These changes can be modelled as a set of differential equations:

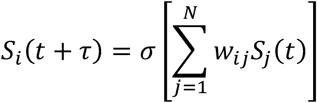

Here, *σ*(*x*) is the sign function similar to Wagner’s model and *w*_*ij*_ is the weight of interaction between gene *i* and gene *j*. The connectivity matrix is given by *w*. Overall connectivity of the network (*c*) is given by the average fraction of entries in *w* that are different from zeros (i.e. *c ∈* (0,1)). The effect of gene duplication on a random network was assessed as in Wagner^34^.

The effect of gene deletion on a random network was assessed as follows. A network of size *N* was considered. The initial and final states of the gene network were randomly chosen where each individual gene has an equal probability of being on or off 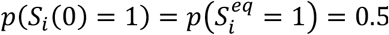. Entries in *w* were randomly and independently chosen from the Gaussian distribution (μ = 0, *σ* = 1). Only weight matrices that lead to stable expression patterns 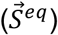 as *t* → ∞ were retained. We iterated the differential equation 100 times to confirm if *w* results in 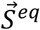 when the initial state is 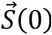. If not, a new *w* was chosen. Once the network set 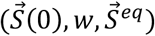 was found, *k* genes (*k ∈* (1,9)) were randomly deleted from the 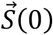 to give 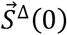. This was done by randomly setting the expression state of the *k* genes to 0. The connectivity matrix *w* was updated such that the effect of remaining genes were set to 0. However, the effect of the deleted genes on the remaining genes were randomly selected from the Gaussian distribution.

The updated connectivity matrix *w* was then used to simulate the new final expression state 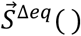. Hamming distance (*d*_*h*_) was calculated between 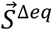 and 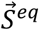. For each *k*, 1000 iterations of simulation were done. The effect of gene deletion was reported as the proportion of instances where *d*_*h*_ > *k*. Calculation of proportion of novel stable networks for gene gain and loss was performed using custom scripts.

## Acknowledgments

This material is based upon work supported by the National Science Foundation under Grant Nos. MCB-1715894 and MCB-1412738. We wish to acknowledge Professors Hisayoshi Nozaki and Takashi Hamaji for kindly sharing *Pandorina morum* and valuable data with us, Hunter Goddard for providing insight on custom scripts, and Professors Stephen Miller, Christopher Toomajian, and Stephanie Shames for their feedback on early drafts of the manuscript. Special thanks to Dan Andresen, Kyle Hutson, Dave Turner for support on the BeoCat Research Cluster at Kansas State University, and XSEDE for super computing support.

## Footnotes

a Hereafter referred to by genus

