## Supplemental Figures and Tables for "Gene loss during a transition to multicellularity"

**This PDF file includes:**

Figs. S1 to S16
Tables S1 to S9

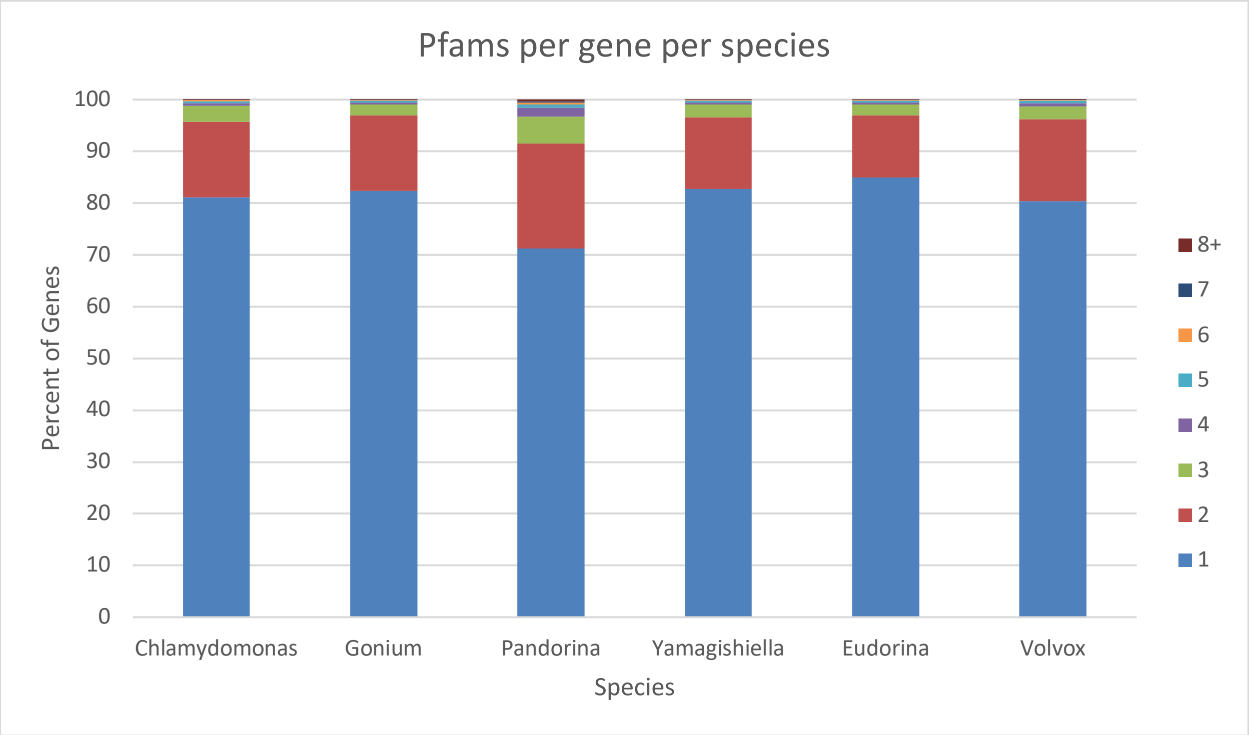


**Figure S1.** Number of Pfams per gene does not change in the five volvocine algae species.

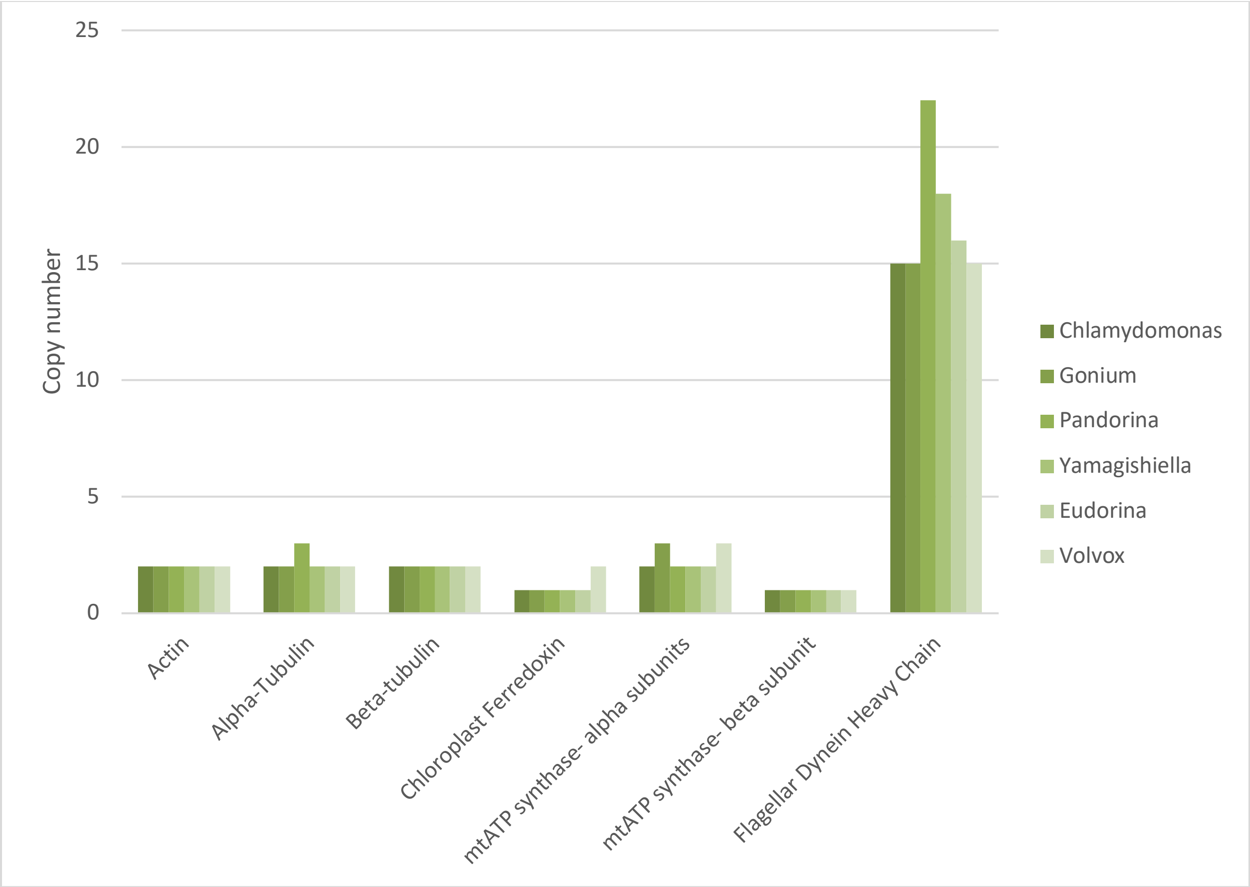


**Figure S2.** Gene copy numbers of nine internal control genes in the six volvocine species.

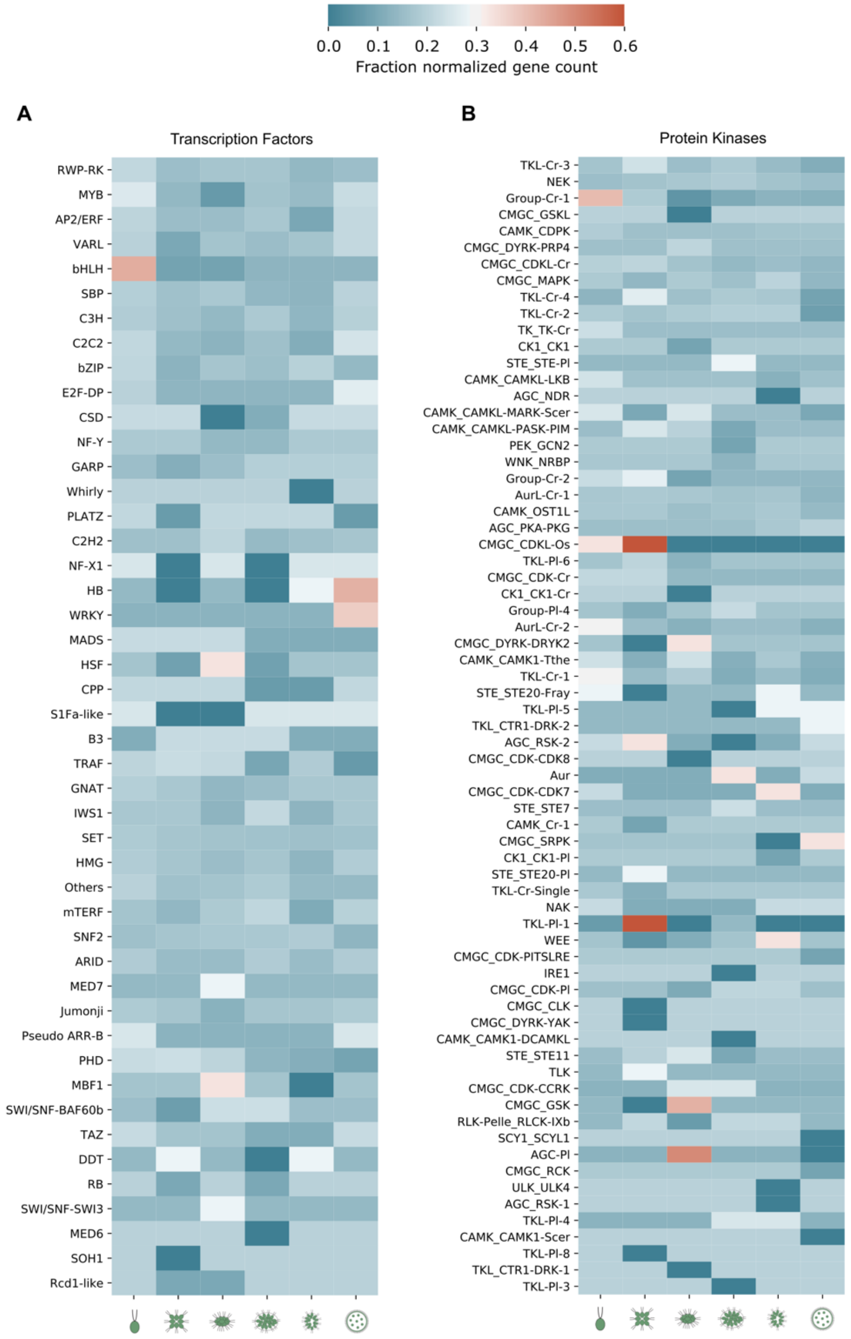


**Figure S3.** (A) Transcription factor and (B) protein kinase family loss and gain in the six volvocine genomes. Relative gene counts represent the fraction normalized count relative to counts of the other species.

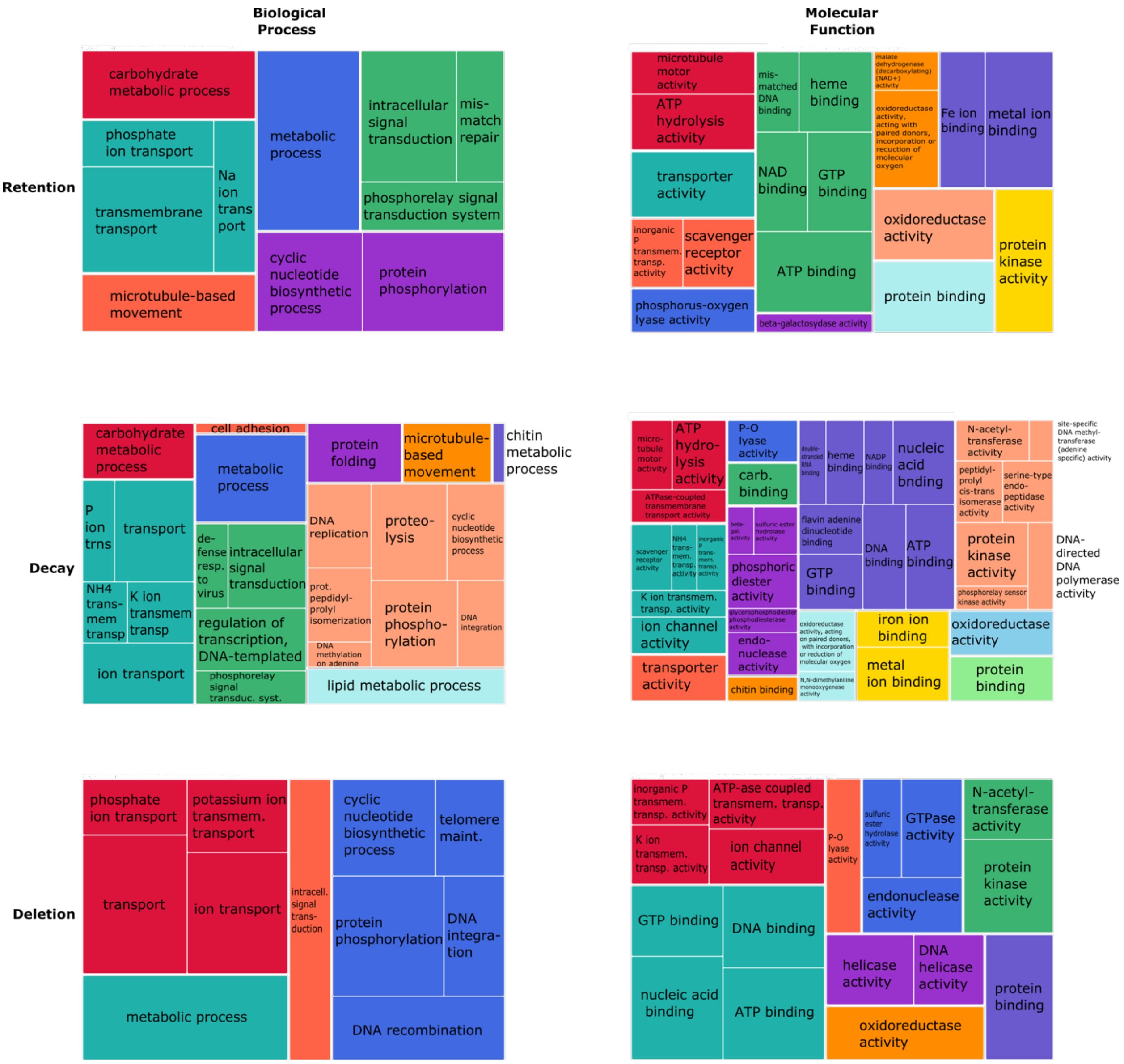


**Figure S4.** GO term representation for biological process and molecular functions of *Chlamydomonas* orthologs of retained (55.8% of total retained genes), decayed (24% of total decayed genes), and deleted (25.3% of total deleted genes) genes.

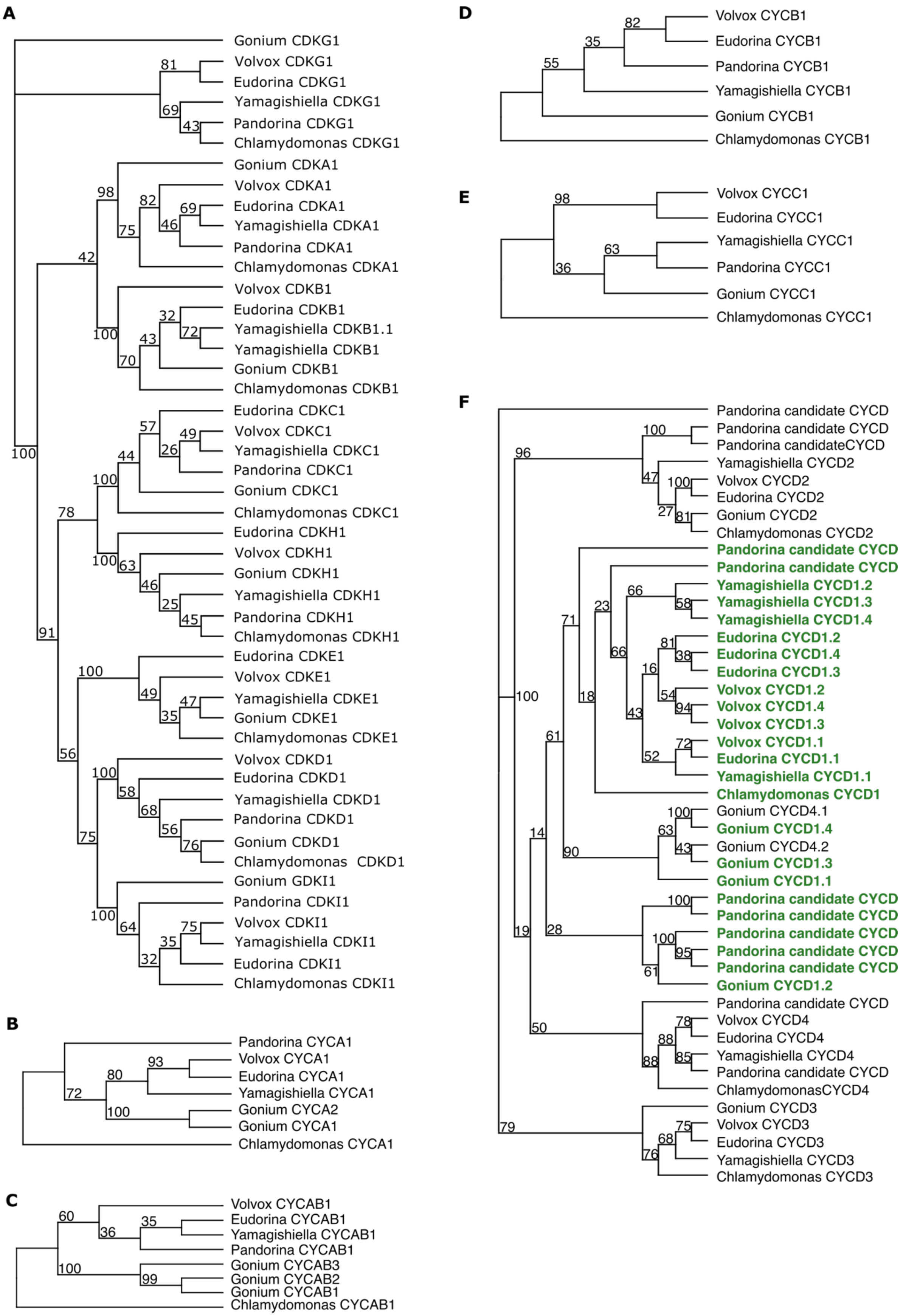
A
Gonium CDKG1
Volvox CDKG1
Eudorina CDKG1
Yamagishiella CDKG1
Pandorina CDKG1
Chlamydomonas CDKG1
Gonium CDKA1
Volvox CDKA1
Eudorina CDKA1
Yamagishiella CDKA1
Pandorina CDKA1
Chlamydomonas CDKA1
Volvox CDKB1
Eudorina CDKB1
Yamagishiella CDKB1.1
Yamagishiella CDKB1
Gonium CDKB1
Chlamydomonas CDKB1
Eudorina CDKC1
Volvox CDKC1
Yamagishiella CDKC1
Pandorina CDKC1
Gonium CDKC1
Chlamydomonas CDKC1
Eudorina CDKH1
Volvox CDKH1
Gonium CDKH1
Yamagishiella CDKH1
Pandorina CDKH1
Chlamydomonas CDKH1
Eudorina CDKE1
Volvox CDKE1
Yamagishiella CDKE1
Gonium CDKE1
Chlamydomonas CDKE1
Volvox CDKD1
Eudorina CDKD1
Yamagishiella CDKD1
Pandorina CDKD1
Gonium CDKD1
Chlamydomonas CDKD1
Gonium GDKI1
Pandorina CDKI1
Volvox CDKI1
Yamagishiella CDKI1
Eudorina CDKI1
Chlamydomonas CDKI1
B
Pandorina CYCA1
Volvox CYCA1
Eudorina CYCA1
Yamagishiella CYCA1
Gonium CYCA2
Gonium CYCA1
Chlamydomonas CYCA1
C
Volvox CYCAB1
Eudorina CYCAB1
Yamagishiella CYCAB1
Pandorina CYCAB1
Gonium CYCAB3
Gonium CYCAB2
Gonium CYCAB1
Chlamydomonas CYCAB1
D
Volvox CYCB1
Eudorina CYCB1
Pandorina CYCB1
Yamagishiella CYCB1
Gonium CYCB1
Chlamydomonas CYCB1
E
Volvox CYCC1
Eudorina CYCC1
Yamagishiella CYCC1
Pandorina CYCC1
Gonium CYCC1
Chlamydomonas CYCC1
F
Pandorina candidate CYCD
Pandorina candidate CYCD
Pandorina candidateCYCD
Yamagishiella CYCD2
Volvox CYCD2
Eudorina CYCD2
Gonium CYCD2
Chlamydomonas CYCD2
Pandorina candidate CYCD
Pandorina candidate CYCD
Yamagishiella CYCD1.2
Yamagishiella CYCD1.3
Yamagishiella CYCD1.4
Eudorina CYCD1.2
Eudorina CYCD1.4
Eudorina CYCD1.3
Volvox CYCD1.2
Volvox CYCD1.4
Volvox CYCD1.3
Volvox CYCD1.1
Eudorina CYCD1.1
Yamagishiella CYCD1.1
Chlamydomonas CYCD1
Gonium CYCD4.1
Gonium CYCD1.4
Gonium CYCD4.2
Gonium CYCD1.3
Gonium CYCD1.1
Pandorina candidate CYCD
Pandorina candidate CYCD
Pandorina candidate CYCD
Pandorina candidate CYCD
Pandorina candidate CYCD
Gonium CYCD1.2
Pandorina candidate CYCD
Volvox CYCD4
Eudorina CYCD4
Yamagishiella CYCD4
Pandorina candidate CYCD
ChlamydomonasCYCD4
Gonium CYCD3
Volvox CYCD3
Eudorina CYCD3
Yamagishiella CYCD3
Chlamydomonas CYCD3

**Figure S5.** Phylogenetic analysis of cell cycle genes. (A) Phylogeny of cyclin dependent kinases (CDKs). (B-F) Phylogeny of cyclins (cyc). CycD1s are in green.

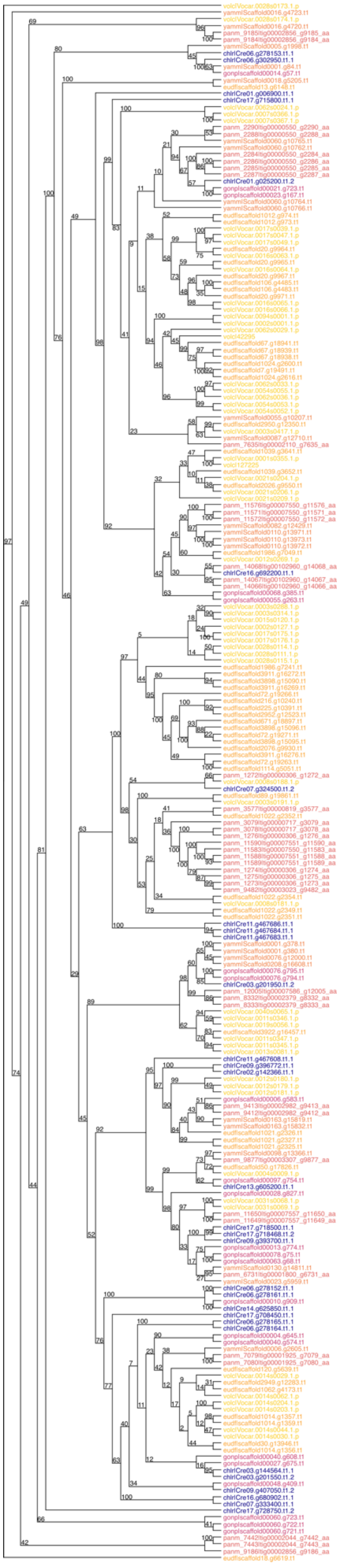

**Figure S6.** Phylogenetic analysis of matrix metalloprotease (MMP) genes. *Chlamydomonas* genes are in blue, *Gonium* in magenta, *Yamagishiella* in salmon, *Eudorina* in orange, and *Volvox* in yellow. Values on the nodes represent percentage bootstrap support.

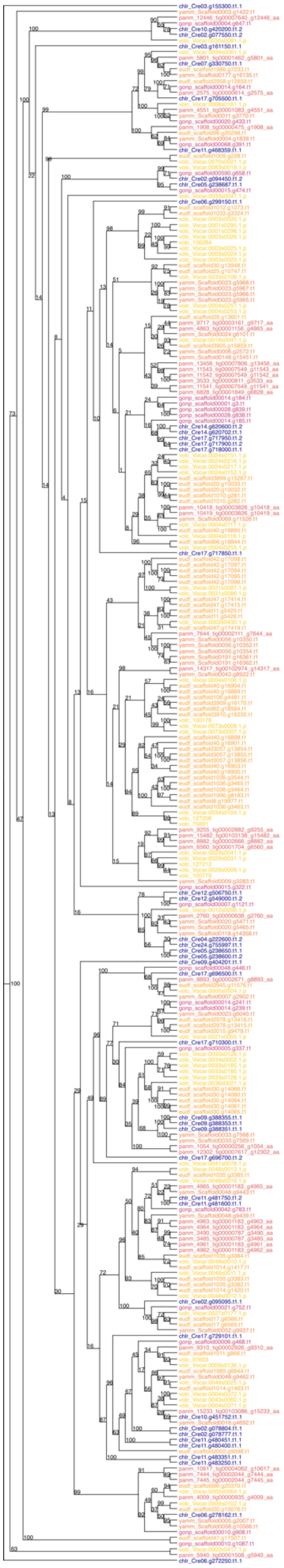

**Figure S7.** Phylogenetic analysis of pherophorin genes. *Chlamydomonas* genes are in blue, *Gonium* in magenta, *Yamagishiella* in salmon, *Eudorina* in orange, and *Volvox* in yellow. Values on the nodes represent percentage bootstrap support.

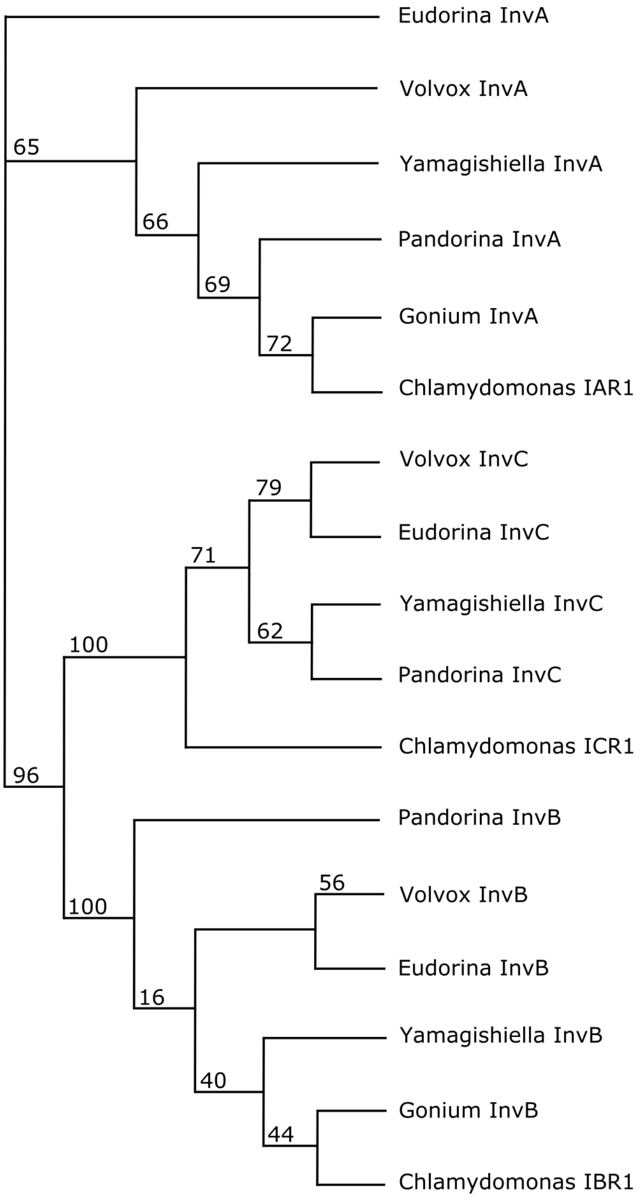


**Figure S8.** Phylogenetic analysis of *inv* genes. Values on the nodes represent percentage bootstrap support.

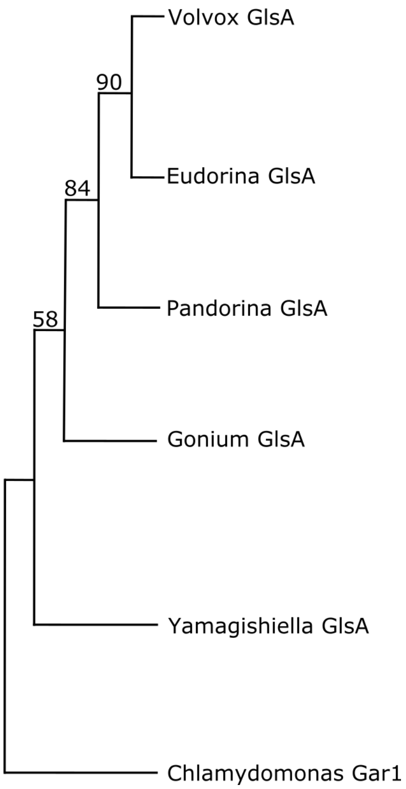


**Figure S9.** Phylogenetic analysis of *glsA* genes. Values on the nodes represent percentage bootstrap support.

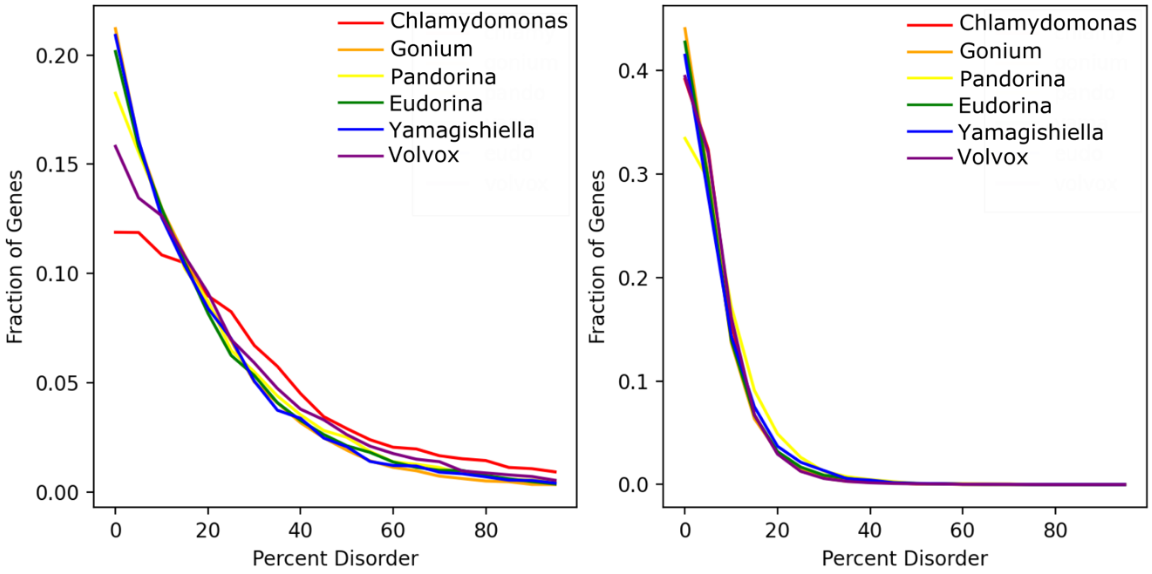


**Figure S10.** Percentage frequency distribution of disordered proteins in the volvocine algae. **A**, Percentage distribution of genes encoding protein-binding proteins with an increasing amount of disorder. **B**, Percentage distribution of genes encoding proteins with an increasing amount of disorder.

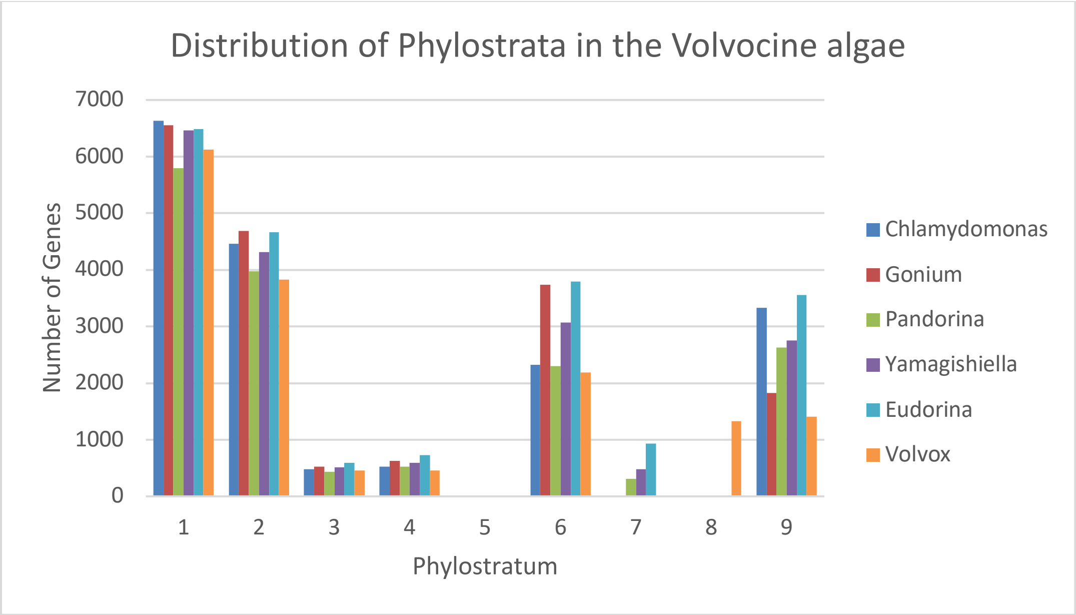


**Figure S11.** Number of genes per phylostratum (PS) for six volvocine species. PS-1 corresponds to cellular organisms (least specific) and PS-9 corresponds to species (most specific).

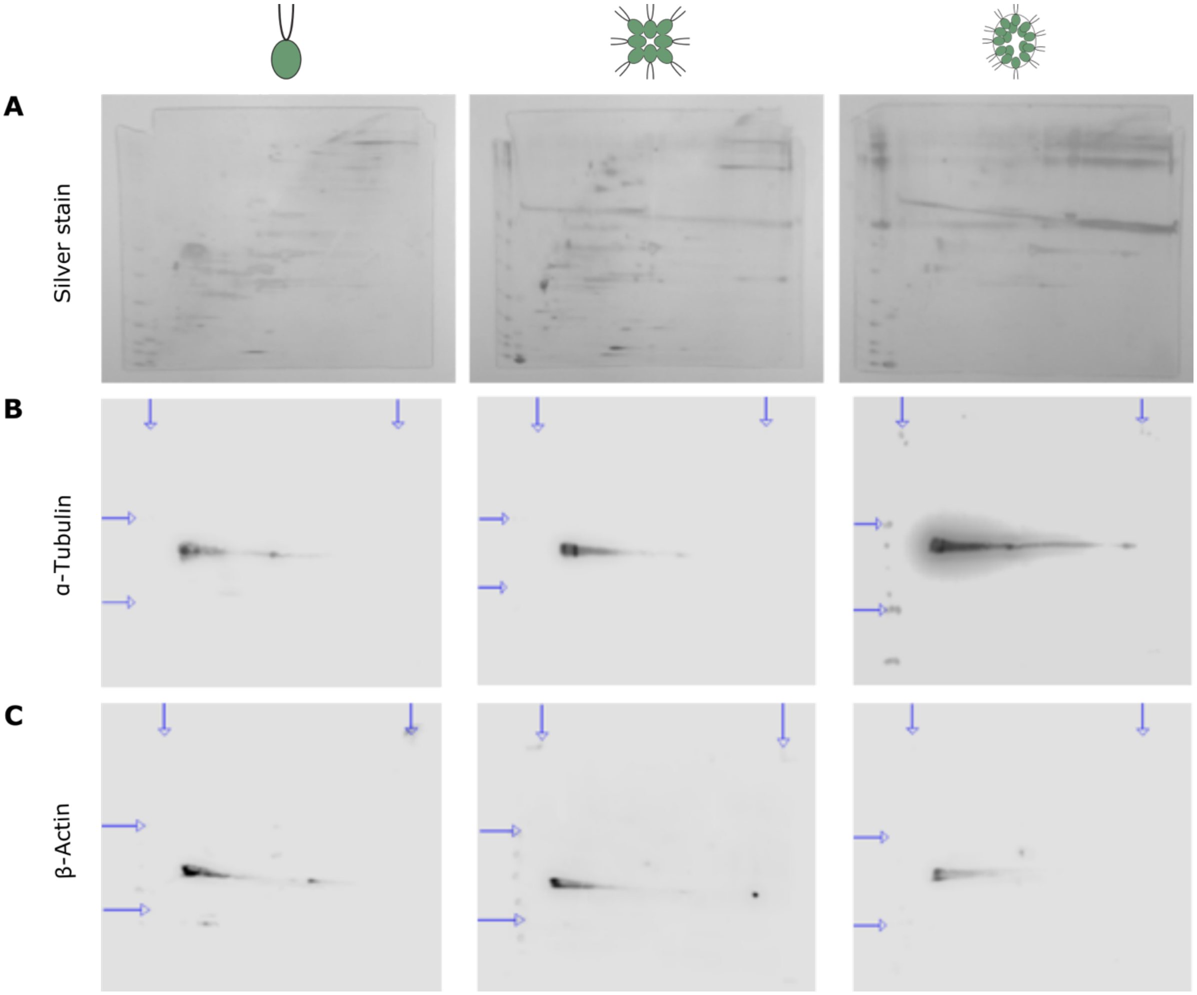


**Figure S12.** Raw images for (A) Silver stained gels; (B) α-tubulin and (C) β-actin blots for *Chlamydomonas, Gonium* and *Eudorina*. Vertical blue arrows indicate boundaries of the native gel slice. Horizontal blue arrows indicate 80 kDa (top) and 25 kDa (bottom) bands on the SDS-PAGE standards ladder. Silver stains are notched at the top corners to indicate the boundaries of the native gel slice.

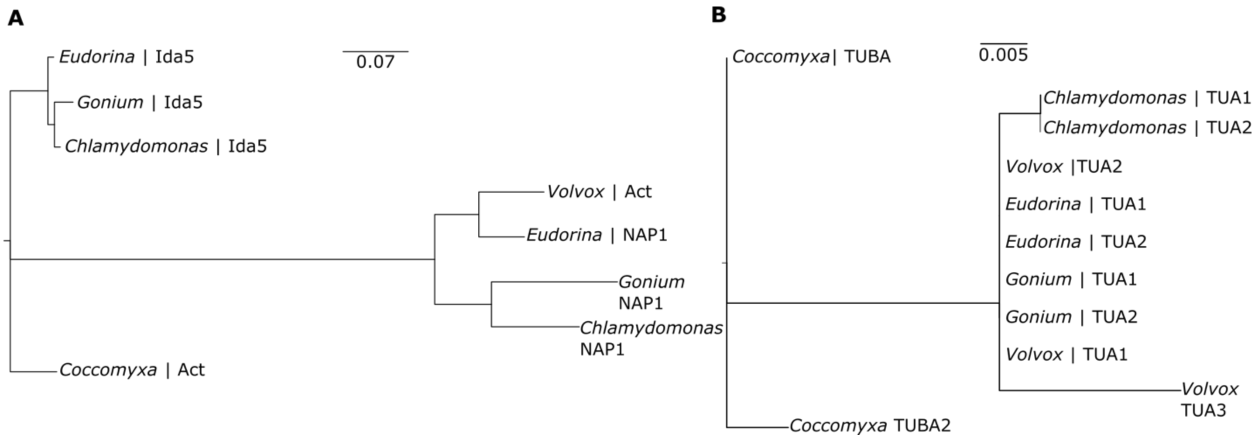


**Figure S13.** Phylogenetic analysis of (A) β-actin and (B) α-tubulin using *Coccomyxa subellipsoidea* (Chlorophyta) as an outgroup.

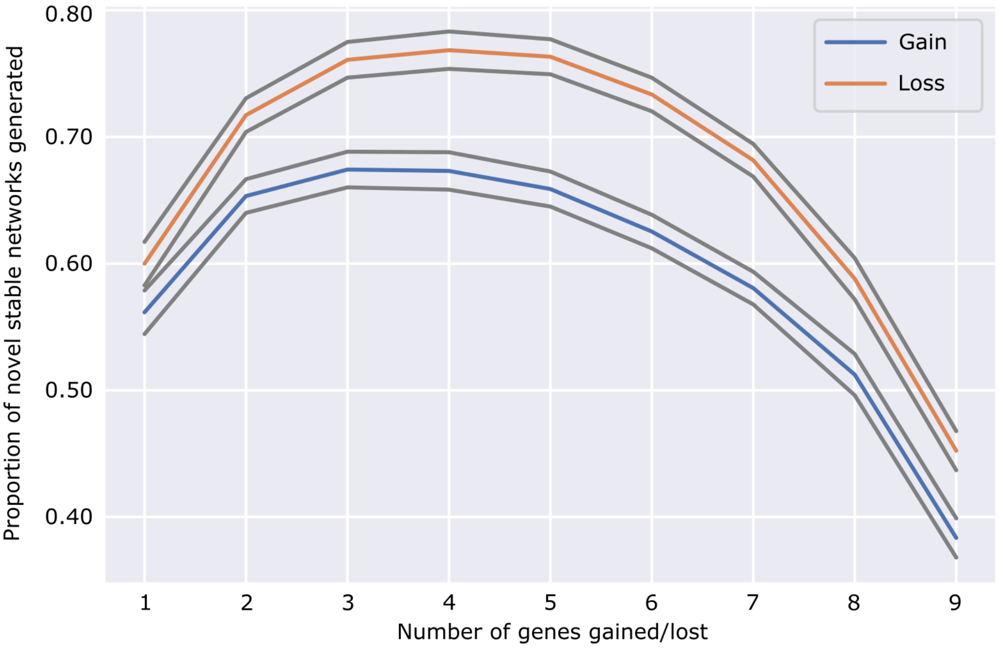


**Figure S14.** Wagner model simulates loss (orange) and duplication (blue) of *k* genes out of a network of *N*=10 genes over different network connectivities. Gene loss yields a higher proportion of novel stable network states than gene gain regardless of changes to network connectivity. Grey lines represent variance.

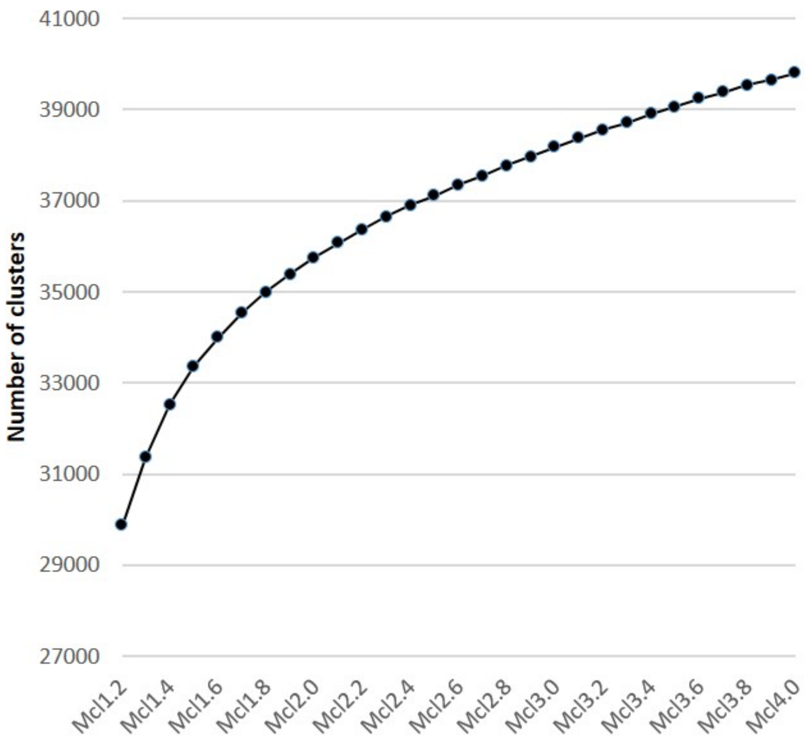


**Figure S15.** Number of orthoMCL clusters for a range of inflation values. Singletons not included.

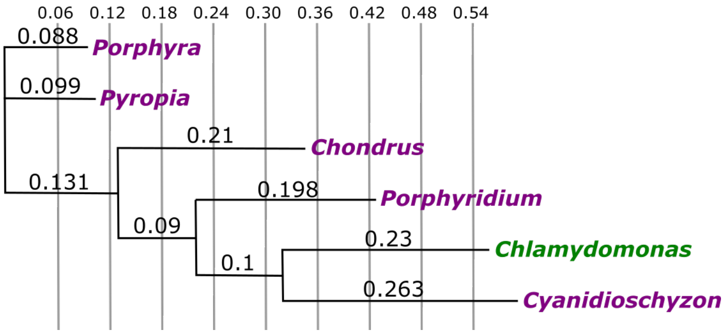


**Figure S16.** Genome-wide phylogeny of red algae (magenta) compared to a chlorophycean outgroup (green) inferred by PosiGene. Branch values indicate distance from the nearest node.

**Table S1.** Genomic features of five volvocine algae.

| | *Chlamydomonas* | *Gonium* | *Pandorina* | *Yamagishiella* Minus | *Yamagishiella* Plus | *Eudorina* Plus | *Eudorina* Minus | *Volvox* |
|---|---|---|---|---|---|---|---|---|
| % G and C content | 64.1 | 64.5 | 62.8 | 62.9 | 62.8 | 63.4 | 63.2 | 56 |
| Protein coding Genes | 17741 | 17948 | 15976 | 18180 | 18416 | 20744 | 22924 | 14247 |
| Total transcripts | 19526 | 17984 | 16542 | 30755 | 31705 | 32233 | 38492 | 16075 |
| Ave Exons Per transcript | 8.62 | 6.94 | 8.67 | 7.06 | 6.93 | 6.38 | 5.73 | 7.91 |
| Average Exon length | 260.99 | 211.04 | 215.8 | 205.93 | 205.37 | 207.78 | 211.58 | 254.39 |
| Average Intron length | 279.17 | 408.89 | 457.7 | 348.37 | 348.28 | 396.73 | 382.3 | 399.5 |
| Average UTR5 length | 203.5 | 232.57 | 187.64 | 235.97 | 237.61 | 235.23 | 233.72 | 231.69 |
| Average UTR3 length | 772.74 | 711.23 | 426.37 | 702.19 | 708.88 | 725.83 | 769.03 | 1105.09 |

**Table S2.** BUSCO completeness values of five volvocine algae annotated genomes.

| Species | Complete (%) | Single (%) | Duplicated (%) | Fragmented (%) | Missing (%) | Total |
|---|---|---|---|---|---|---|
| *Chlamydomonas* | 1503 (99) | 1411 (92.9) | 92 (6.1) | 5 (0.3) | 11 (0.7) | 1519 |
| *Gonium* | 1415 (93.2) | 1400 (92.2) | 15 (1) | 69 (4.5) | 35 (2.3) | 1519 |
| *Pandorina* | 1193 (78.5) | 1115 (73.4) | 78 (5.1) | 46 (3) | 280 (18.5) | 1519 |
| *Yamagishiella* | 1422 (93.6) | 992 (65.3) | 430 (28.3) | 58 (3.8) | 39 (2.6) | 1519 |
| *Eudorina* | 1340 (88.2) | 838 (55.2) | 502 (33) | 106 (7) | 73 (4.8) | 1519 |
| *Volvox* | 1464 (96.4) | 1332 (87.7) | 132 (8.7) | 16 (1.1) | 39 (2.5) | 1519 |

Completeness values describe how many BUSCOS are conserved between query genomes and the Chlorophyta database. BUSCOS are classified as Complete, Fragmented, and Missing. Complete BUSCOS can be Single or Duplicated.

**Table S3.** Orthologous group and Pfam domain counts for six volvocine species. Species names are encoded as follows: G=*Gonium,* P=*Pandorina*, Y=*Yamagishiella*, E=*Eudorina*, V=*Volvox*

| Present in | Number of orthologous groups | Number of Pfams |
|---|---|---|
| C | 473 | 12 |
| G | 453 | 15 |
| P | 537 | 18 |
| Y | 317 | 11 |
| E | 411 | 17 |
| V | 226 | 23 |
| CG | 161 | 7 |
| CP | 50 | 4 |
| CY | 103 | 2 |
| CE | 91 | 1 |
| CV | 80 | 6 |
| GP | 78 | 2 |
| GY | 162 | 3 |
| GE | 120 | 5 |
| GV | 46 | 1 |
| PY | 98 | 0 |
| PE | 67 | 4 |
| PV | 33 | 0 |
| YE | 254 | 3 |
| YV | 96 | 1 |
| EV | 255 | 6 |
| CGP | 36 | 1 |
| CGY | 82 | 4 |
| CGE | 45 | 2 |
| CGV | 39 | 6 |
| CPY | 32 | 2 |
| CPE | 14 | 1 |
| CPV | 20 | 2 |
| CYE | 51 | 1 |
| CYV | 47 | 5 |
| CEV | 58 | 0 |
| GPY | 47 | 3 |
| GPE | 39 | 0 |
| GPV | 15 | 1 |
| GYE | 86 | 3 |

| | | |
|---|---|---|
| GYV | 40 | 1 |
| GEV | 46 | 1 |
| PYE | 66 | 1 |
| PYV | 34 | 1 |
| PEV | 50 | 0 |
| YEV | 129 | 1 |
| CGPY | 68 | 7 |
| CGPE | 44 | 3 |
| CGPV | 26 | 12 |
| CGYE | 132 | 5 |
| CGYV | 72 | 12 |
| CGEV | 79 | 9 |
| CPYE | 84 | 1 |
| CPYV | 35 | 12 |
| CPEV | 38 | 6 |
| CYEV | 164 | 8 |
| GPYE | 88 | 3 |
| GPYV | 33 | 2 |
| GPEV | 37 | 2 |
| GYEV | 122 | 2 |
| PYEV | 124 | 3 |
| CGPYE | 354 | 31 |
| CGPYV | 145 | 68 |
| CGPEV | 120 | 43 |
| CGYEV | 1352 | 91 |
| CPYEV | 442 | 52 |
| GPYEV | 212 | 10 |
| CGPYEV | 6297 | 1184 |
| Total | 15155 | 1743 |

**Table S4.** Conserved gene counts for six volvocine species.

| Species | Actin | mtATP-A | mtATP-B | Flagellar Dynein heavy chain | Chloroplast Ferridoxin | α-Tubulin | β-Tubulin | β-2 Tubulin |
|---|---|---|---|---|---|---|---|---|
| *Chlamydomonas* | 2 | 2 | 1 | 15 | 1 | 2 | 2 | 3 |
| *Gonium* | 2 | 3 | 1 | 15 | 1 | 2 | 2 | 4 |
| *Pandorina* | 2 | 2 | 1 | 22 | 1 | 3 | 2 | |
| *Yamagishiella* | 2 | 2 | 1 | 18 | 1 | 2 | 2 | 2 |
| *Eudorina* | 2 | 2 | 1 | 16 | 1 | 2 | 2 | 2 |
| *Volvox* | 2 | 3 | 1 | 15 | 2 | 2 | 2 | 2 |

**Table S5**. Histone H1 gene counts for volvocine algae.

| Species | H1 |
|---|---|
| *Chlamydomonas* | 3 |
| *Gonium* | 3 |
| *Pandorina* | 2 |
| *Yamagishiella* | 3 |
| *Eudorina* | 2 |
| *Volvox* | 4 |

**Table S6.** Histone family copy number and tail variant number of five volvocine algae.

| Histone | Species | Copy number | Tail var. number |
|---|---|---|---|
| H2A | *Chlamydomonas* | 31 | 5 |
| | *Gonium* | 34 | 5 |
| | *Pandorina* | 30 | 1 |
| | *Yamagishiella* | 19 | 3 |
| | *Eudorina* | 19 | 4 |
| | *Volvox* | 15 | 3 |
| H2B | *Chlamydomonas* | 29 | 9 |
| | *Gonium* | 35 | 12 |
| | *Pandorina* | 32 | 9 |
| | *Yamagishiella* | 20 | 13 |
| | *Eudorina* | 18 | 14 |
| | *Volvox* | 15 | 12 |
| H3 | *Chlamydomonas* | 35 | 4 |
| | *Gonium* | 30 | 24 |
| | *Pandorina* | 30 | 1 |
| | *Yamagishiella* | 21 | 6 |
| | *Eudorina* | 23 | 4 |
| | *Volvox* | 13 | 3 |
| H4 | *Chlamydomonas* | 32 | 2 |
| | *Gonium* | 34 | 3 |
| | *Pandorina* | 30 | 1 |
| | *Yamagishiella* | 27 | 3 |
| | *Eudorina* | 21 | 2 |
| | *Volvox* | 14 | 2 |

**Table S7.** Histone gene counts for Rhodophytes.

| Species | H1 | H2A | H2B | H3 | H4 |
|---|---|---|---|---|---|
| *Porphyra* | 1 | 6 | 4 | 74 | 2 |
| *Pyropia* | 1 | 3 | 1 | 22 | 1 |
| *Chondrus* | 1 | 3 | 2 | 21 | 3 |
| *Porphyridium* | 1 | 3 | 2 | 3 | 2 |
| *Cyanidioschyzon* | 1 | 3 | 2 | 2 | 2 |

**Table S8.** Distribution and type of gene losses in multicellular volvocine algae.

| Absence in | Total | Decay | Deletion |
|---|---|---|---|
| GPYEV | 1017 | 880 | 137 |
| GPYE | 0 | 0 | 0 |
| GPYV | 8 | 6 | 2 |
| GPEV | 2 | 1 | 1 |
| GYEV | 5 | 5 | 0 |
| PYEV | 30 | 26 | 4 |
| GPY | 0 | 0 | 0 |
| GPE | 0 | 0 | 0 |
| GPV | 2 | 2 | 0 |
| GYE | 1 | 1 | 0 |
| GYV | 0 | 0 | 0 |
| GEV | 1 | 1 | 0 |
| PYE | 1 | 1 | 0 |
| PYV | 8 | 5 | 3 |
| PEV | 6 | 4 | 2 |
| YEV | 8 | 6 | 2 |
| GP | 1 | 1 | 0 |
| GY | 0 | 0 | 0 |
| PY | 0 | 0 | 0 |
| PE | 2 | 1 | 1 |
| PV | 6 | 6 | 0 |
| GE | 0 | 0 | 0 |
| YE | 0 | 0 | 0 |
| GV | 2 | 1 | 1 |
| EV | 11 | 11 | 0 |
| YV | 5 | 3 | 2 |
| G | 3 | 3 | 0 |
| P | 4 | 3 | 1 |
| Y | 3 | 2 | 1 |
| E | 1 | 1 | 0 |
| V | 14 | 5 | 9 |

Species names are encoded as follows: G=*Gonium,* P=*Pandorina*, Y=*Yamagishiella*, E=*Eudorina*, V=*Volvox.* All values have a P-value<0.01 and reject uniform distribution of decay and deletion events.

**Table S9.** Matrix metalloprotease (MMP) and pherophorin gene counts for five volvocine species.

| Species | MMP genes | Pherophorin genes |
|---|---|---|
| *Chlamydomonas* | 34 | 41 |
| *Gonium* | 22 | 21 |
| *Pandorina* | 42 | 38 |
| *Yamagishiella* | 38 | 32 |
| *Eudorina* | 63 | 69 |
| *Volvox* | 58 | 68 |
